# MOVICS: an R package for multi-omics integration and visualization in cancer subtyping

**DOI:** 10.1101/2020.09.15.297820

**Authors:** Xiaofan Lu, Jialin Meng, Yujie Zhou, Liyun Jiang, Fangrong Yan

**Affiliations:** State Key Laboratory of Natural Medicines, Research Center of Biostatistics and Computational Pharmacy, China Pharmaceutical University, Nanjing 210009, China; Department of Urology, The First Affiliated Hospital of Anhui Medical University; Institute of Urology & Anhui Province Key Laboratory of Genitourinary Diseases, Anhui Medical University, Hefei, Anhui 230022, China; Division of Gastroenterology and Hepatology, Key Laboratory of Gastroenterology and Hepatology, Ministry of Health, Shanghai Institute of Digestive Disease, Renji Hospital, School of Medicine, Shanghai Jiao Tong University, Shanghai 200001, China; Department of Biostatistics, The University of Texas MD Anderson Cancer Center, Texas 77030, USA

## Abstract

**Summary:** Stratification of cancer patients into distinct molecular subgroups based on multi-omics data is an important issue in the context of precision medicine. Here we present *MOVICS*, an R package for multi-omics integration and visualization in cancer subtyping. *MOVICS* provides a unified interface for 10 state-of-the-art multi-omics integrative clustering algorithms, and incorporates the most commonly used downstream analyses in cancer subtyping researches, including characterization and comparison of identified subtypes from multiple perspectives, and verification of subtypes in external cohort using a model-free approach for multiclass prediction. *MOVICS* also creates feature rich customizable visualizations with minimal effort.

**Availability and implementation:** *MOVICS* package and online tutorial are freely available at https://github.com/xlucpu/MOVICS.

## 1 Introduction

Recent advances in next-generation sequencing, microarrays and mass spectrometry for omics data production have enabled the generation and collection of different modalities of high-dimensional molecular data. Combining these various types of multi-omics datasets could have the potential to reveal further systems-level insights, thus enhancing a comprehensive view of the mechanisms of disease or biological process.

So far, more than a dozen multi-omics integrative clustering algorithms based on different programming languages have been published and applied in medical research (Rappoport and Shamir, 2018). Although these algorithms have rigorous theoretical derivation and practical applications, most of them only considered whether the subtypes can show distinct prognosis. Moreover, the lack of unified input and output of different algorithms makes it cumbersome to concatenate downstream analyses. To our best knowledge, *CancerSubtypes* (Xu, et al., 2017) and *CEPICS* (Duan, et al., 2019) are two R packages for cancer subtyping studies. They have unified the format of inputs and outputs, but only several algorithms were integrated and neither one of them provides function for systematically characterizing subtypes, which is precisely an essential issue in cancer biology and a crucial step towards personalized treatment practices. These two platforms spontaneously avoided somatic mutations which may be the key to cancer heterogeneity. However, compared with continuous variables (i.e., RNA expression, DNA methylation), binary data makes the program more complicated in both data pre-processing and visualization. Moreover, pipeline is scarce regarding identification of subtype-specific biomarkers and verification of subtypes in external cohort, and such analytical procedure is critical to the reproducibility of subtypes.

We herein present the *MOVICS* R package, an integrated analytic pipeline which provides a unified interface for 10 state-of-the-art multi-omics clustering algorithms, most of which have been evaluated for performance according to the literature (Pierre-Jean, et al., 2019). Particularly, *MOVICS* incorporates the most commonly used downstream analyses in cancer subtyping researches, including the characterization and comparison of identified subtypes from multiple perspectives (*e.g*., prognosis, somatic mutation, fraction genome altered, drug sensitivity, partitions agreement), and verification of subtypes in external cohort using a model-free approach for multiclass prediction. *MOVICS* also creates feature rich customizable visualizations with minimal effort. We documented the detailed usage and analysis process of *MOVICS* in the Supplementary Material.

With the explosion of biological data generation, *MOVICS* will serve a wide range of users, including biologists, bioinformaticians and computational biologists, in analysing cancer subtypes from data, and assist cancer therapy by moving away from the ‘one-size-fits-all’ approach to patient care.

## 2 Implementation and main functions

*MOVICS* currently requires R (≥4.0.0) and mainly comprises three modules (*i.e*., GET, COMP and RUN). All user-oriented functions are named starting from the label of any module (Fig. 1).

**Fig. 1.**
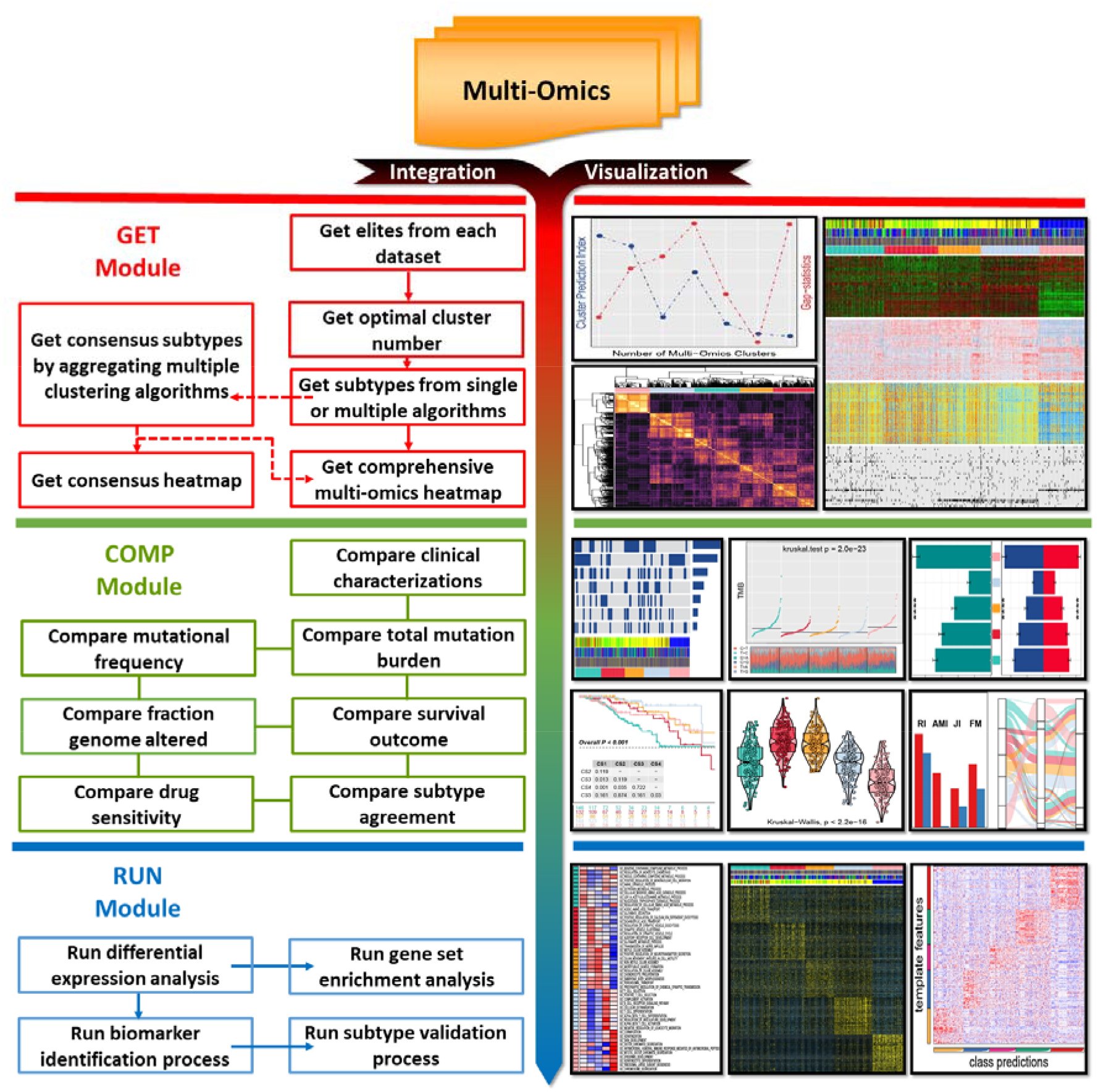
Schematic view of *MOVICS* pipeline for integration and visualization of multi-omics data. The left panel describes three built-in analytic modules. The directed arrow indicates that the function should be executed sequentially, while the undirected solid line indicates that the functions are independent of each other, and the optional functions are connected by a dotted line. The right panel presents the simplified illustrations generated by *MOVICS*.

### 2.1 GET Module

Using GET functions, users can filter out a portion of elites for each omics data separately based on four methods: standard deviation (*sd*), median absolute deviation (*mad*), mutational frequency (*freq*), and univariate Cox proportional hazards regression (*cox*). *MOVICS* then refers to clustering prediction index (Chalise and Fridley, 2017) and Gaps-statistics (Hastie, et al., 2001) to estimate the number of clusters. Clustering number at which the sum of the two statistics reaches the maximum will be arbitrarily considered optimal, but we suggests to reconsider the number with reference to prior knowledge of molecular subtypes of the given cancer type. Further in this module, *MOVICS* provides a unified interface for users to specify one out of ten or multiple multi-omics clustering algorithms simultaneously. Borrowing the idea of consensus ensembles which is a later integration approach for aggregating results from multiple algorithms (Strehl and Ghosh, 2002), *MOVICS* calculates a consensus matrix which represents a robust pairwise similarities for samples and applies supervised clustering to derive stable subtypes. All clustering results, whether based on a single algorithm or multiple algorithms, maintain the same outputs to facilitate downstream analyses, along with a vertically connected multi-omics heatmap obtained to visualize the genome-wide patterns within or among subtypes.

### 2.2 COMP Module

Within this module, *MOVICS* provides the following seven functions for users to realize the most commonly used downstream analyses in cancer subtyping. 1) Users can evaluate prognosis to see how well the survival patterns are discriminated from different subtypes. 2) Clinical features can be compared and also summarized with a formatted .docx file that is easy to use in medical research papers. 3) For genetic analyse, mutational frequency can be compared to infer the subtype-specific driver mutations. 4) We added functionality to allow users to predict patients’ response to drugs (*e.g*., chemotherapies) by estimating IC_50_ individually (Geeleher, et al., 2014). Since immunotherapy is becoming a pillar of modern cancer treatment, two functions are established to calculate and compare 5) total mutation burden (Chan, et al., 2019) and 6) fraction genome altered (Davoli, et al., 2017) that may influence immune response. 7) Additionally, *MOVICS* calculates four statistics: Rand Index, Adjusted Mutual Information, Jaccard Index, and Fowlkes-Mallows to evaluate and visualize the agreement of new subtypes with previous classifications, which is critical to reflect the robustness of clustering analysis and to determine potential but novel subtypes.

### 2.3 RUN Module

This module runs the identification process to determine the subtype-specific biomarkers and functional pathways, and runs validation procedure for subtype reproducibility by starting with pairwise differential expression analyses. *MOVICS* embeds three mainstream methods, namely, edgeR (Robinson, et al., 2010), DESeq2 (Love, et al., 2014), and limma (Ritchie, et al., 2015), which can adapt to various expression profiles (*e.g*., RNA-Seq count, microarrays expression profile, and normalized expression matrix). *MOVICS* performs gene set enrichment analysis to retrieve a functional profile of each subtype in order to better understand the underlying biological processes (Yu, et al., 2012). To verify the current subtypes in external cohort, *MOVICS* first identifies a certain number of unique biomarkers for each subtype based on log_2_FoldChange to build a template, and then determines the most likely subtype of each sample by harnessing nearest template prediction (Hoshida, 2010).

## 3 Application and conclusion

We detailed the use of *MOVICS* in the Supplementary Materials by a complete analysis process with two breast cancer cohorts (TCGA-BRCA as training and Yau-BRCA as validation). All generated charts meet the quality requirements of publication and can be further edited locally.

In summary, *MOVICS* provides a suite for multi-omics integration and visualization in cancer subtyping.

## Funding

This work was supported by the National Key R&D Program of China (2019YFC1711000), the National Natural Science Foundation of China (81973145), and the “Double First-Class” University Project (CPU2018GY09).

### Conflict of interest

*none declared*

## Acknowledgement

We would like to express our gratitude to *Dr. Morgane Pierre-Jean* for the inspiration brought by the study of evaluating unsupervised methods for multi-omics data integration. We also want to thank *Dr. Enyu Lin* for the helping in calculation and visualization of fraction genome altered, and to thank *Dr. Rongfang Shen* for the assistance in visualization of Transitions and Transversions. At last, sincere thanks to the brilliant contributors of all the functions incorporated in *MOVICS* package.

## Notes

https://github.com/xlucpu/MOVICS

### Competing Interest Statement

The authors have declared no competing interest.

