## Supplementary material for "MOVICS: an R package for multi-omics integration and visualization in cancer subtyping"

#### Contents

**1 Introduction** **2**

**2 Installation** **2**

**3 Real Data Scenario** **2**

**4 MOVICS Pipeline** **3**

**5 Little Trick** **38**

**6 Summary** **40**

**7 References** **40**

*Correspondence to: State Key Laboratory of Natural Medicines, Research Center of Biostatistics and Computational Pharmacy, China Pharmaceutical University, Nanjing 210009, China.

### 1 Introduction

Recent advances in next-generation sequencing, microarrays and mass spectrometry for omics data production have enabled the generation and collection of different modalities of high-dimensional molecular data[1]. Clustering multi-omic data has the potential to reveal further systems-level insights, but raises computational and biological challenges[2]. We aim to show in this supplementary material how to use MOVICS to perform multi-omics integrative clustering and visualization for cancer subtyping researches. This R package provides a unified interface for 10 state-of-the-art multi-omics clustering algorithms, and standardizes the output for each algorithm so as to form a pipeline for downstream analyses. Ten algorithms are CIMLR[3], iClusterBayes[4], MoCluster[5], COCA[6], ConsensusClustering[7], IntNMF[8], LRAcluster[9], NEMO[10], PINSPlus[11], and SNF[12] where the former three methods can also perform the process of feature selection. For cancer subtyping studies, MOVICS also forms a pipeline for most commonly used downstream analyses for further subtype characterization and creates editable publication-quality illustrations.

### 2 Installation

It is essential that you have R 4.0.2[13] or above already installed on your computer or server. MOVICS is a pipeline that utilizes many other R packages that are currently available from CRAN and Bioconductor. For all of the steps of the pipeline to work, make sure that you have upgraded Bioconductor to newest version (BiocManager v3.11). After you have R and Bioconductor installed properly, type the following code into your R session:

```
if (!requireNamespace("BiocManager", quietly = TRUE))
    install.packages("BiocManager")
if (!require("devtools"))
    install.packages("devtools")
devtools::install_github("xlucpu/MOVICS", host = "https://api.github.com")
```

When you are installing MOVICS, you may encounter some errors saying that some packages are not installed. These errors are caused by recursively depending on R packages, so if one package was not installed properly on your computer, MOVICS would fail. To solve these errors, you simply need to check those error messages, find out which packages required are missing, then install it with command `BiocManager::install("YourErrorPackage")` or `install.packages("YourErrorPackage")` directly. After that, retry installing MOVICS, it may take several times, but eventually it should work. Or, We highly suggest that you referred to the `Imports` in the DESCRIPTION file, try to install all the R dependencies, and then install MOVICS. After installation, you should be able to load the MOVICS package in your R session:

```
library("MOVICS")
```

### 3 Real Data Scenario

This package contains two pre-processed real datasets of breast cancer. One dataset is `brca.tcga.RData` which is a list that includes 643 samples with four complete omics data types of breast cancer retrieved from TCGA-BRCA cohort[14] (*i.e.*, mRNA expression, lncRNA expression, DNA methylation profiling and somatic mutation matrix), and corresponding clinicopathological information (*i.e.*, age, pathological stage, PAM50 subtype, vital status and overall suvival time); such data list also contains corresponding RNA-Seq raw count table and Fragments Per Kilobase Million (FPKM) data in order to test the functions for downstream analyses (*e.g.*, differential expression analysis, drug sensitivity analysis, *etc*). The other one, `brca.yau.RData`, is an external validation dataset which contains gene expression profiles and clinicopathological information that downloaded from BRCA-YAU cohort[15] with 682 samples (one sample without annotation of PAM50 subtype was removed), which can be used to test the predictive functions available in MOVICS. These two datasets can be loaded like below:

```
# load example data of breast cancer
load(system.file("extdata", "brca.tcga.RData", package = "MOVICS", mustWork = TRUE))
load(system.file("extdata", "brca.yau.RData",  package = "MOVICS", mustWork = TRUE))
```

Since in most cases, multi-omics clustering procedure is rather time-consuming, the TCGA-BRCA dataset was therefore first pre-processed to extract top 500 mRNAs, 500 lncRNA, 1,000 promoter CGI probes/genes with high variation using statistics of median absolute deviation (MAD), and 30 genes that mutated in at least 3% of the entire cohort. All these features are used in the following clustering analyses.

### 4 MOVICS Pipeline

#### 4.1 Pipeline Introduction

MOVICS Pipeline comprises separate functions that can be categorized into three modules using three colors:

- **GET Module: get subtypes through multi-omics integrative clustering**
  - `getElites()`: get elites which are those features that pass the filtering procedure and are used for analyses
  - `getClustNum()`: get optimal cluster number by calculating clustering prediction index (CPI) and Gap-statistics
  - `get%algorithm_name%()`: get results from one specific multi-omics integrative clustering algorithm with detailed parameters
  - `getMOIC()`: get a list of results from multiple multi-omics integrative clustering algorithm with parameters by default
  - `getConsensusMOIC()`: get a consensus matrix that indicates the clustering robustness across different clustering algorithms and generate a consensus heatmap
  - `getStdiz()`: get a standardized data for generating comprehensive multi-omics heatmap
  - `getMoHeatmap()`: get a comprehensive multi-omics heatmap based on clustering results
- **COMP Module: compare subtypes from multiple perspectives**
  - `compSurv()`: compare survival outcome and generate a Kalan-Meier curve with pairwise comparison if possible
  - `compClinvar()`: compare and summarize clinical features among different identified subtypes
  - `compMut()`: compare mutational frequency and generate an OncoPrint with significant mutations
  - `compTMB()`: compare total mutation burden among subtypes and generate distribution of Transitions and Transversions
  - `compFGA()`: compare fraction genome altered among subtypes and generate a barplot for distribution comparison
  - `compDrugsen()`: compare estimated half maximal inhibitory concentration ($IC_{50}$) for drug sensitivity and generate a boxviolin for distribution comparison
  - `compAgree()`: compare agreement of current subtypes with other pre-existed classifications and generate an alluvial diagram and an agreement barplot
- **RUN Module: run marker identification and verify subtypes**
  - `runDEA()`: run differential expression analysis with three popular methods for choosing, including edgeR, DESeq2, and limma
  - `runMarker()`: run biomarker identification to determine uniquely and significantly differential expressed genes for each subtype
  - `runGSEA()`: run gene set enrichment analysis (GSEA), calculate activity of functional pathways and generate a pathway-specific heatmap
  - `runNTP()`: run nearest template prediction based on identified biomarkers to evaluate subtypes in external cohorts

#### 4.2 Steps of Pipeline

Basically, the above three connected modules explain the workflow of this R package. MOVICS first identifies the cancer subtype (CS) by using one or multiple clustering algorithms; if multiple clustering algorithms are specified, it is highly recommended to perform a consensus clustering based on different subtyping results in order to derive stable and robust subtypes. Second, after having subtypes it is natural to exploit the heterogeneity of subtypes from as many angles as possible. Third, each subtype should have a list of subtype-specific biomarkers for reproducibility in external cohorts.

##### 4.2.1 GET Module

**4.2.1.1 get data from example files** We first extracted from the list of `brca.tcga`, including 4 types of multi-omics data with exactly the same order for samples. Except for omics data, this list also contains RNA-Seq raw count, FPKM matrix, MAF data, segmented copy number, clinical and survival information for downstream analyses.

```
# print name of example data
names(brca.tcga)
#> [1] "mRNA.expr"   "lncRNA.expr" "meth.beta"   "mut.status"  "count"
#> [6] "fpkm"        "maf"         "segment"     "clin.info"
names(brca.yau)
#> [1] "mRNA.expr" "clin.info"

# extract multi-omics data
mo.data   <- brca.tcga[1:4]

# extract raw count data for downstream analyses
count     <- brca.tcga$count

# extract fpkm data for downstream analyses
fpkm      <- brca.tcga$fpkm

# extract maf for downstream analysis
maf       <- brca.tcga$maf

# extract segmented copy number for downstream analyses
segment   <- brca.tcga$segment

# extract survival information
surv.info <- brca.tcga$clin.info
```

**4.2.1.2 get elites by reducing data dimension** Although all these omics data have been already processed (filtered from the original dataset), we here to show how to use `getElites()` function to filter out features that meet some stringent requirements, and those features that are preserved in this procedure are considered elites by MOVICS. Four filtering methods are provided, namely `mad` for median absolute deviation, `sd` for standard deviation, `cox` for univariate Cox proportional hazards regression, and `freq` for binary omics data. This function also handles missing values coded in $NA$ by removing them directly or imputing them by $k$ nearest neighbors using a Euclidean metric[16] through argument of `na.action`.

```
# scenario 1:
# considering we are dealing with an expression data that have 2 rows with NA values
tmp       <- brca.tcga$mRNA.expr # get expression data
dim(tmp) # check data dimension
#> [1] 500 643
tmp[1,1]  <- tmp[2,2] <- NA # set 2 rows with NA values
tmp[1:3,1:3] # check data
#>         BRCA-A03L-01A BRCA-A04R-01A BRCA-A075-01A
#> SCGB2A2            NA          1.42          7.24
#> SCGB1D2         10.11            NA          5.88
#> PIP              4.54          2.59          4.35
elite.tmp <- getElites(dat       = tmp,
                       method    = "mad",
```

```
                       na.action = "rm", # NA values will be removed
                       elite.pct = 1) # all (100%) features will be selected
#> --2 features with NA values are removed.
#> missing elite.num then use elite.pct
dim(elite.tmp$elite.dat) # we have removed 2 rows with NA data
#> [1] 498 643

elite.tmp <- getElites(dat       = tmp,
                       method    = "mad",
                       na.action = "impute", # NA values will be imputed
                       elite.pct = 1)
#> missing elite.num then use elite.pct
dim(elite.tmp$elite.dat) # all data kept
#> [1] 500 643
elite.tmp$elite.dat[1:3,1:3] # NA values have been imputed
#>         BRCA-A03L-01A BRCA-A04R-01A BRCA-A075-01A
#> SCGB2A2         6.867         1.420          7.24
#> SCGB1D2        10.110         4.739          5.88
#> PIP             4.540         2.590          4.35

# scenario 2:
# considering we are dealing with continuous data and use mad or sd to select elites
tmp       <- brca.tcga$mRNA.expr # get expression data with 500 features
elite.tmp <- getElites(dat       = tmp,
                       method    = "mad",
                       elite.pct = 0.1) # top 10% features are kept
#> missing elite.num then use elite.pct
dim(elite.tmp$elite.dat) # get 50 elite left
#> [1]  50 643

elite.tmp <- getElites(dat       = tmp,
                       method    = "sd",
                       elite.num = 100, # top 100 features are kept
                       elite.pct = 0.1) # disabled due to indicated elite.num
#> elite.num has been provided then discards elite.pct.
dim(elite.tmp$elite.dat) # get 100 elites left
#> [1] 100 643

# scenario 3:
# considering we are dealing with data and use cox to select elite
tmp       <- brca.tcga$mRNA.expr # get expression data
elite.tmp <- getElites(dat       = tmp,
                       method    = "cox",
                       surv.info = surv.info, # survival information
                       p.cutoff  = 0.05,
                       elite.num = 100) # disabled because cox refers to p.cutoff
#> --all sample matched between omics matrix and survival data.
#> 5% 10% 15% 20% 25% 30% 35% 40% 45% 50% 55% 60% 65% 70% 75% 80% 85% 90% 95% 100%
dim(elite.tmp$elite.dat) # get 125 elites
#> [1] 125 643
table(elite.tmp$unicox$pvalue < 0.05) # 125 genes have pvalue<0.05 in Cox
#>
#> FALSE  TRUE
#>   375   125
```

```
tmp       <- brca.tcga$mut.status # get mutation data
elite.tmp <- getElites(dat       = tmp,
                       method    = "cox",
                       surv.info = surv.info,
                       p.cutoff  = 0.05,
                       elite.num = 100)
#> --all sample matched between omics matrix and survival data.
#> 7% 13% 20% 27% 33% 40% 47% 53% 60% 67% 73% 80% 87% 93% 100%
dim(elite.tmp$elite.dat) # get 3 elites
#> [1]   3 643
table(elite.tmp$unicox$pvalue < 0.05) # 3 mutations have pvalue<0.05
#>
#> FALSE  TRUE
#>    27     3

# scenario 4:
# considering we are dealing with mutation data using freq to select elites
tmp       <- brca.tcga$mut.status # get mutation data
rowSums(tmp) # check mutation frequency
#> PIK3CA   TP53    TTN   CDH1  GATA3   MLL3  MUC16 MAP3K1  SYNE1  MUC12    DMD
#>    208    186    111     83     58     49     48     38     33     32     31
#>  NCOR1    FLG   PTEN   RYR2  USH2A  SPTA1 MAP2K4  MUC5B    NEB   SPEN  MACF1
#>     31     30     29     27     27     25     25     24     24     23     23
#>   RYR3    DST  HUWE1  HMCN1  CSMD1  OBSCN   APOB  SYNE2
#>     23     22     22     22     21     21     21     21
elite.tmp <- getElites(dat       = tmp,
                       method    = "freq", # must set as 'freq'
                       elite.num = 80, # now refer to frequency of mutation
                       elite.pct = 0.1) # disabled due to indicated elite.num
#> --method of 'freq' only supports binary omics data (e.g., somatic mutation matrix), and in this manner
#> elite.num has been provided then discards elite.pct.
rowSums(elite.tmp$elite.dat) # genes mutated in >80 samples are kept
#> PIK3CA   TP53    TTN   CDH1
#>    208    186    111     83

elite.tmp <- getElites(dat       = tmp,
                       method    = "freq",
                       elite.pct = 0.2) # now refer to mutation/sample size
#> --method of 'freq' only supports binary omics data (e.g., somatic mutation matrix), and in this manner
#> missing elite.num then use elite.pct
rowSums(elite.tmp$elite.dat) # genes mutated in >0.2*643=128.6 samples are kept
#> PIK3CA   TP53
#>    208    186

# get mo.data list just like below (not run)
# mo.data <- list(omics1 = elite.tmp$elite.dat,
#                 omics2 = ...)
```

Since the `mo.data` has been already prepared, We am going to stick on this data list, and take you further to MOVICS.

**4.2.1.3 get optimal number for clustering** The most important parameter to estimate in any clustering study is the optimum number of clusters $k$ for the data, where $k$ needs to be small enough to reduce noise but large enough to retain important information. Herein MOVICS refers to CPI[8] and Gaps-statistics[17] to estimate the number of clusters by using `getClustNum()` function.

```r
# identify optimal clustering number (may take a while)
optk.brca <- getClustNum(data        = mo.data,
                         is.binary   = c(F,F,F,T), # the 4th data is a binary matrix
                         try.N.clust = 2:8, # try cluster number from 2 to 8
                         fig.name    = "CLUSTER NUMBER OF TCGA-BRCA")
#> calculating Cluster Prediction Index...
#> 5% complete
#> 5% complete
#> 10% complete
#> 10% complete
#> 15% complete
#> 15% complete
#> 20% complete
#> 25% complete
#> 25% complete
#> 30% complete
#> 30% complete
#> 35% complete
#> 35% complete
#> 40% complete
#> 45% complete
#> 45% complete
#> 50% complete
#> 50% complete
#> 55% complete
#> 55% complete
#> 60% complete
#> 65% complete
#> 65% complete
#> 70% complete
#> 70% complete
#> 75% complete
#> 75% complete
#> 80% complete
#> 85% complete
#> 85% complete
#> 90% complete
#> 90% complete
#> 95% complete
#> 95% complete
#> 100% complete
#> calculating Gap-statistics...
#> visualization done...
#> --the imputed optimal cluster number is 3 arbitrarily, but it would be better referring to other prior
```

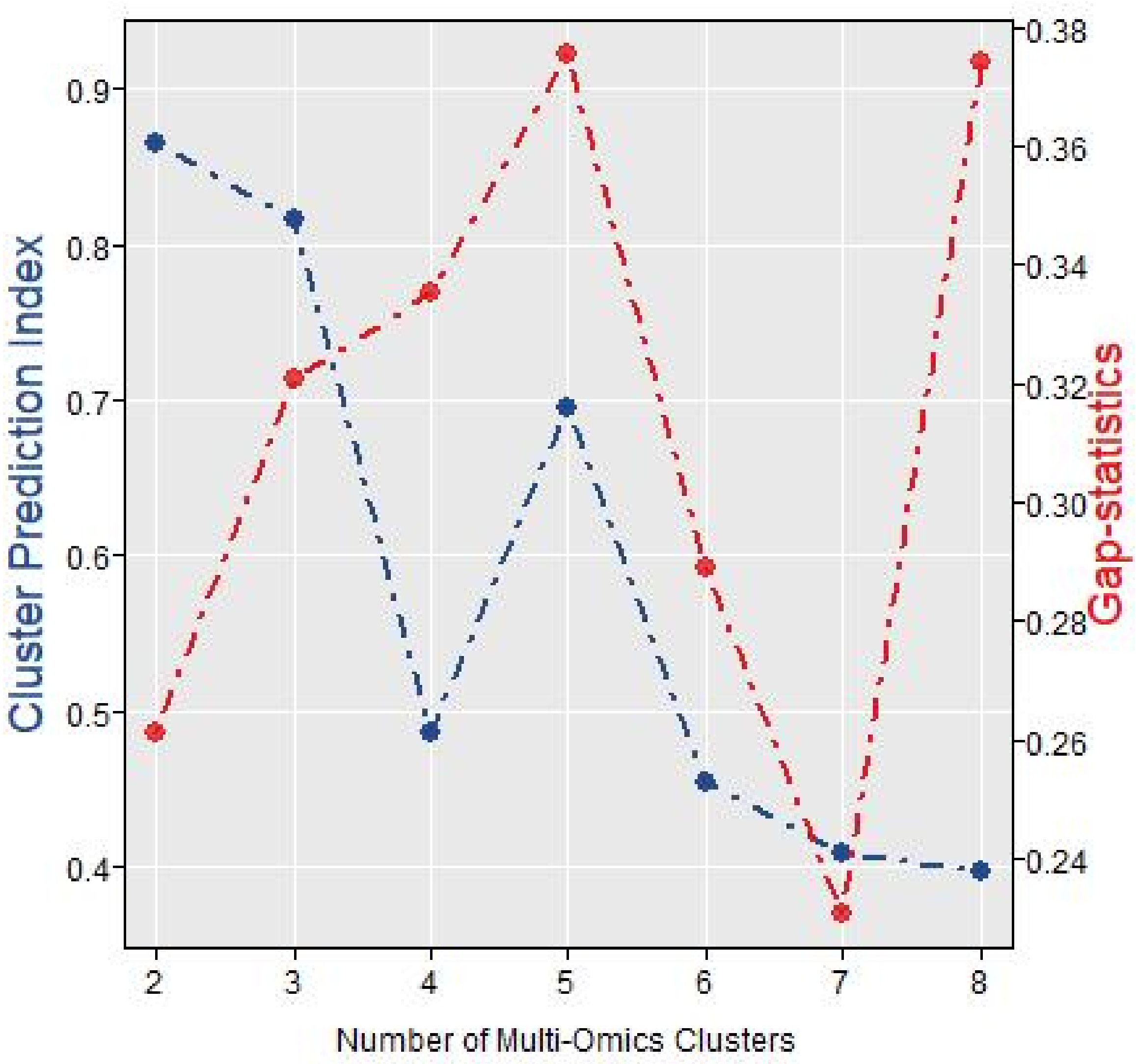

Figure 1: Identification of optimal cluster number by calculating CPI (blue line) and Gaps-statistics (red line) in TCGA-BRCA cohort.

The above estimation of clustering number gives an arbitrary *k* of 3. However, the popular PAM50 classifier for breast cancer has 5 classifications and if taking a close look at the descriptive figure, both CPI and Gaps-statistics does not decline too much at *k* of 5. Therefore, *k* of 5 is chosen as the optimal clustering number for further analyses.

**4.2.1.4 get results from single algorithm** In this part we first use MOVICS to perform multi-omics integrative clustering by specifying one algorithm (*i.e.*, iClusterBayes) with detailed parameters.

```
# perform iClusterBayes (may take a while)
iClusterBayes.res <-
  getiClusterBayes(data        = mo.data,
                   N.clust     = 5,
                   type        = c("gaussian","gaussian","gaussian","binomial"),
                   n.burnin    = 1800,
                   n.draw      = 1200,
                   prior.gamma = c(0.5, 0.5, 0.5, 0.5),
                   sdev        = 0.05,
                   thin        = 3)
#> clustering done...
#> feature selection done...
```

Otherwise, a unified function can be used for all algorithms individually with detailed parameters, that is, `getMOIC()`.

```
iClusterBayes.res <-
  getMOIC(data        = mo.data,
          N.clust     = 5,
          methodslist = "iClusterBayes", # specify only ONE algorithm here
          type        = c("gaussian","gaussian","gaussian","binomial"), # data type
          n.burnin    = 1800,
          n.draw      = 1200,
          prior.gamma = c(0.5, 0.5, 0.5, 0.5),
          sdev        = 0.05,
          thin        = 3)
#> clustering done...
#> feature selection done...
#> iClusterBayes done...
```

By specifying only one algorithm (*i.e.*, iClusterBayes) in the argument of `methodslist`, the above `getMOIC()` will return exactly the same results from `getiClusterBayes()` if the same parameters are provided. The returned result contains a `clust.res` object that has two columns: `clust` to indicate the subtype which the sample belongs to, and `samID` records the corresponding sample name. For algorithms that provide feature selection procedure (*i.e.*, iClusterBayes, CIMLR, and MoCluster), the result also contains a `feat.res` object that stores the information of such procedure. To those algorithms involving hierarchical clustering (*e.g.*, COCA, ConsensusClustering), the corresponding dendrogram for sample clustering will be also returned as `clust.dend`, which is useful if the users want to put them at the heatmap.

**4.2.1.5 get results from multiple algorithms at once** If a list of algorithms to `methodslist` argument are simultaneously specified in `getMOIC()`, it will automatically perform each algorithm with default parameters one by one, and a list of results derived from specified algorithms will be finally returned. Now that iClusterBayes has been finished, we try other 9 algorithms at once with parameters by default.

```
# perform multi-omics integrative clustering with the rest of 9 algorithms
moic.res.list <-
  getMOIC(data        = mo.data,
          methodslist = list("SNF", "PINSPlus", "NEMO",
                             "COCA", "LRAcluster", "ConsensusClustering",
                             "IntNMF", "CIMLR", "MoCluster"),
          N.clust     = 5,
          type        = c("gaussian", "gaussian", "gaussian", "binomial"))
#> --you choose more than 1 algorithm and all of them shall be run with parameters by default.
#> SNF done...
#> Clustering method: kmeans
#> Perturbation method: noise
#> PINSPlus done...
#> NEMO done...
#> COCA done...
#> LRAcluster done...
#> end fraction
#> clustered
#> clustered
#> clustered
#> clustered
#> ConsensusClustering done...
#> IntNMF done...
#> clustering done...
#> feature selection done...
#> CIMLR done...
#> clustering done...
```

```
#> feature selection done...
#> MoCluster done...

# attach iClusterBayes.res as a list to moic.res.list with 9 results already
moic.res.list <- append(moic.res.list,
                        list("iClusterBayes" = iClusterBayes.res))

# save moic.res.list to local path
save(moic.res.list, file = "moic.res.list.rda")
```

**4.2.1.6 get consensus from different algorithms** Since now all the clustering results from 10 algorithms are in hand, MOVICS borrows the idea of consensus ensembles18 for later integration of the clustering results derived from different algorithms, so as to improve the clustering robustness. To be specific, if $t_{max}$ algorithms are specified where $2 \leq t_{max} \leq 10$, `getConsensusMOIC()` calculates a matrix $M_{n \times n}^{(t)}$ per algorithm where $n$ is the number of samples, and $M_{ij}^{(t)} = 1$ only when the sample $i$ and $j$ are clustered in the same subtype, otherwise $M_{ij}^{(t)} = 0$. After get all results from specified algorithms, MOVICS calculates a consensus matrix $CM = \sum_{t=1}^{t_{max}} M^{(t)}$, and $cm_{ij} \in [0, 10]$. Such matrix represents a robust pairwise similarities for samples because it considers different multi-omics integrative clustering algorithms. MOVICS then searches for a stable clustering result by applying hierarchical clustering to $CM$. The simplest way to perform `getConsensusMOIC()` is to pass a list of object returned by multiple `get%algorithm_name%()` or by `getMOIC()` with specific argument of `methodslist`. This can be done like below:

```
cmoic.brca <- getConsensusMOIC(moic.res.list = moic.res.list,
                               fig.name      = "CONSENSUS HEATMAP",
                               distance      = "euclidean",
                               linkage       = "average")
```

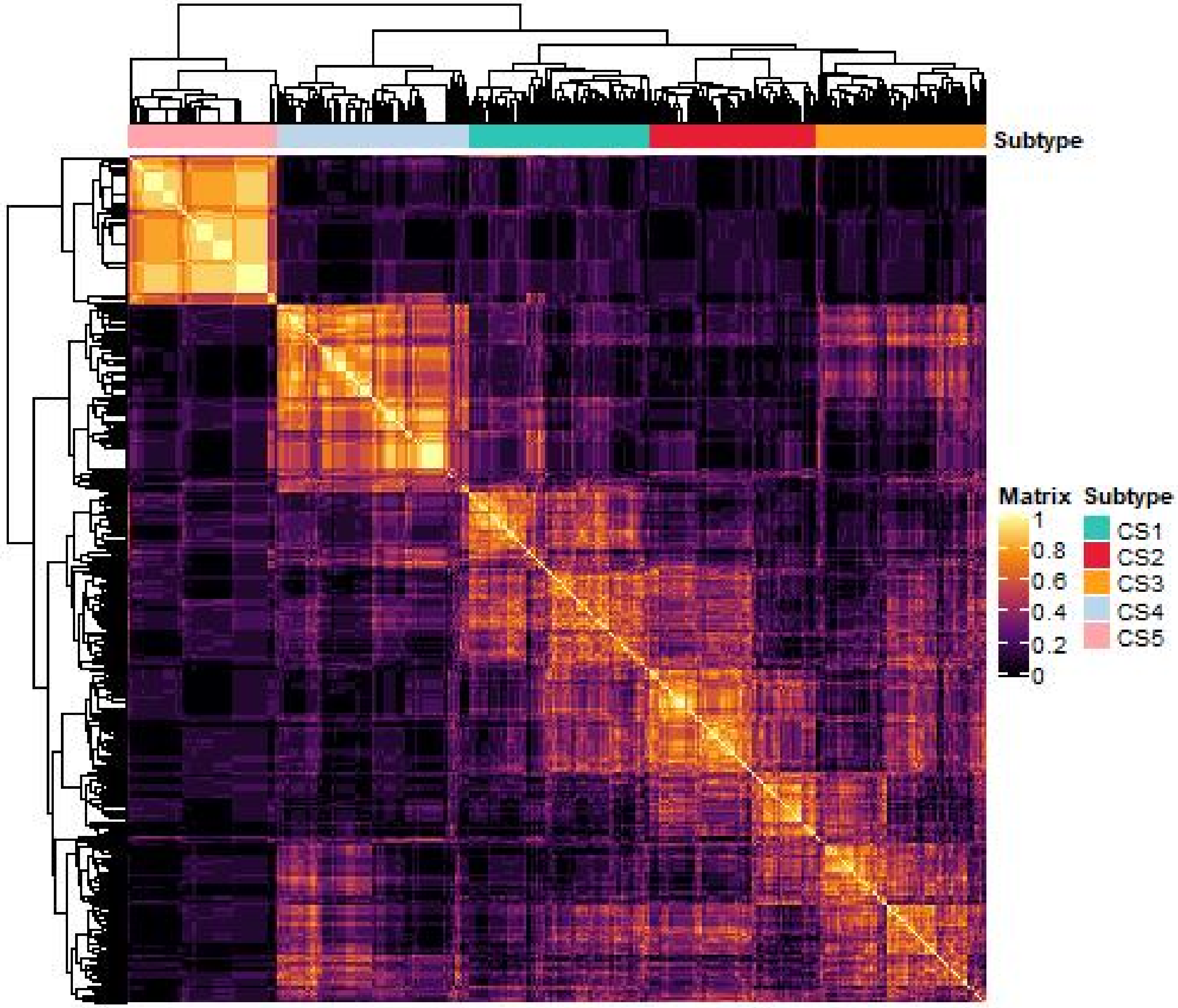

Figure 2: Consensus heatmap based on results from 10 multi-omics integrative clustering algorithms with cluster number of 5.

Remember, you must choose at least two methods to perform the above consensus clustering by `getConsensusMOIC()`, and this function will return a consensus matrix (a probability matrix represents how many times samples belonging to the same subtype can be clustered together by different multi-omics clustering methods) and a corresponding consensus heatmap. Ideally, the consensus heatmap will show a perfect diagonal rectangle, and the input values are 0 and 1 only because all algorithms derived the same clustering results.

**4.2.1.7 get multi-omics heatmap based on clustering result** MOVICS provides `getMoHeatmap` to visually deal with pre-clustered multi-omics data by heatmap. Before using `getMoHeatmap()`, omics data should be properly processed by using the function of `getStdiz()` which returns a list storing normalized omics data. Omics data, especially for expression (*e.g.*, RNA and protein), should be centered (`centerFlag = TRUE`) or scaled (`scaleFlag = TRUE`) or z-scored (both centered and scaled). Generally, DNA methylation data ($\beta$ matrix ranging from 0 to 1) and somatic mutation (0 and 1 binary matrix) should not be normalized. However, it is a good choice to convert the methylation $\beta$ value to M value following the formula of $M = log_2 \frac{\beta}{1-\beta}$ for its stronger signal in visualization and M value is suitable for normalization[19]. This function also provides an argument of `halfwidth` for continuous omics data; such argument is used to truncate the 'extremum' after normalization; specifically, normalized values that exceed the `halfwidth` boundaries will be replaced by the `halfwidth`, which is beneficial to map colors in heatmap.

```r
# convert beta value to M value for stronger signal
indata <- mo.data
indata$meth.beta <- log2(indata$meth.beta / (1 - indata$meth.beta))

# data normalization for heatmap
plotdata <- getStdiz(data       = indata,
                     halfwidth  = c(2,2,2,NA), # no truncation for mutation
                     centerFlag = c(T,T,T,F), # no center for mutation
                     scaleFlag  = c(T,T,T,F)) # no scale for mutation
```

As we mentioned earlier, several algorithms also provide feature selection; those selected features show a complex cross-talk with other omics data and might have special biological significance that drives the heterogeneity of cancers. Therefore we show below how to generate a comprehensive heatmap based on a single algorithm (*e.g.*, iClusterBayes) with selected features. However in the first place, features must be extracted:

```r
feat   <- iClusterBayes.res$feat.res
feat1  <- feat[which(feat$dataset == "mRNA.expr"),][1:10,"feature"]
feat2  <- feat[which(feat$dataset == "lncRNA.expr"),][1:10,"feature"]
feat3  <- feat[which(feat$dataset == "meth.beta"),][1:10,"feature"]
feat4  <- feat[which(feat$dataset == "mut.status"),][1:10,"feature"]
annRow <- list(feat1, feat2, feat3, feat4)
```

The `feat.res` contained in `iClusterBayes.res` is sorted by posterior probability of features for each omics data. In this manner, the top 10 features are selected for each omics data and a feature list is generated and named as `annRow` for heatmap raw annotation.

```r
# set color for each omics data
mRNA.col   <- c("#00FF00", "#008000", "#000000", "#800000", "#FF0000")
lncRNA.col <- c("#6699CC", "white"  , "#FF3C38")
meth.col   <- c("#0074FE", "#96EBF9", "#FEE900", "#F00003")
mut.col    <- c("grey90" , "black")
col.list   <- list(mRNA.col, lncRNA.col, meth.col, mut.col)

# comprehensive heatmap (may take a while)
getMoHeatmap(data          = plotdata,
             row.title     = c("mRNA","lncRNA","Methylation","Mutation"),
             is.binary     = c(F,F,F,T), # the 4th data is a binary matrix
             legend.name   = c("mRNA.FPKM","lncRNA.FPKM","M value","Mutated"),
             clust.res     = iClusterBayes.res$clust.res, # cluster results
             clust.dend    = NULL, # no dendrogram
             show.rownames = c(F,F,F,F), # specify for each omics data
             show.colnames = FALSE, # show no sample names
             annRow        = annRow, # mark selected features
             color         = col.list,
             annCol        = NULL, # no annotation for samples
             annColors     = NULL, # no annotation color
             width         = 10, # width of each subheatmap
             height        = 5, # height of each subheatmap
             fig.name      = "COMPREHENSIVE HEATMAP OF ICLUSTERBAYES")
```

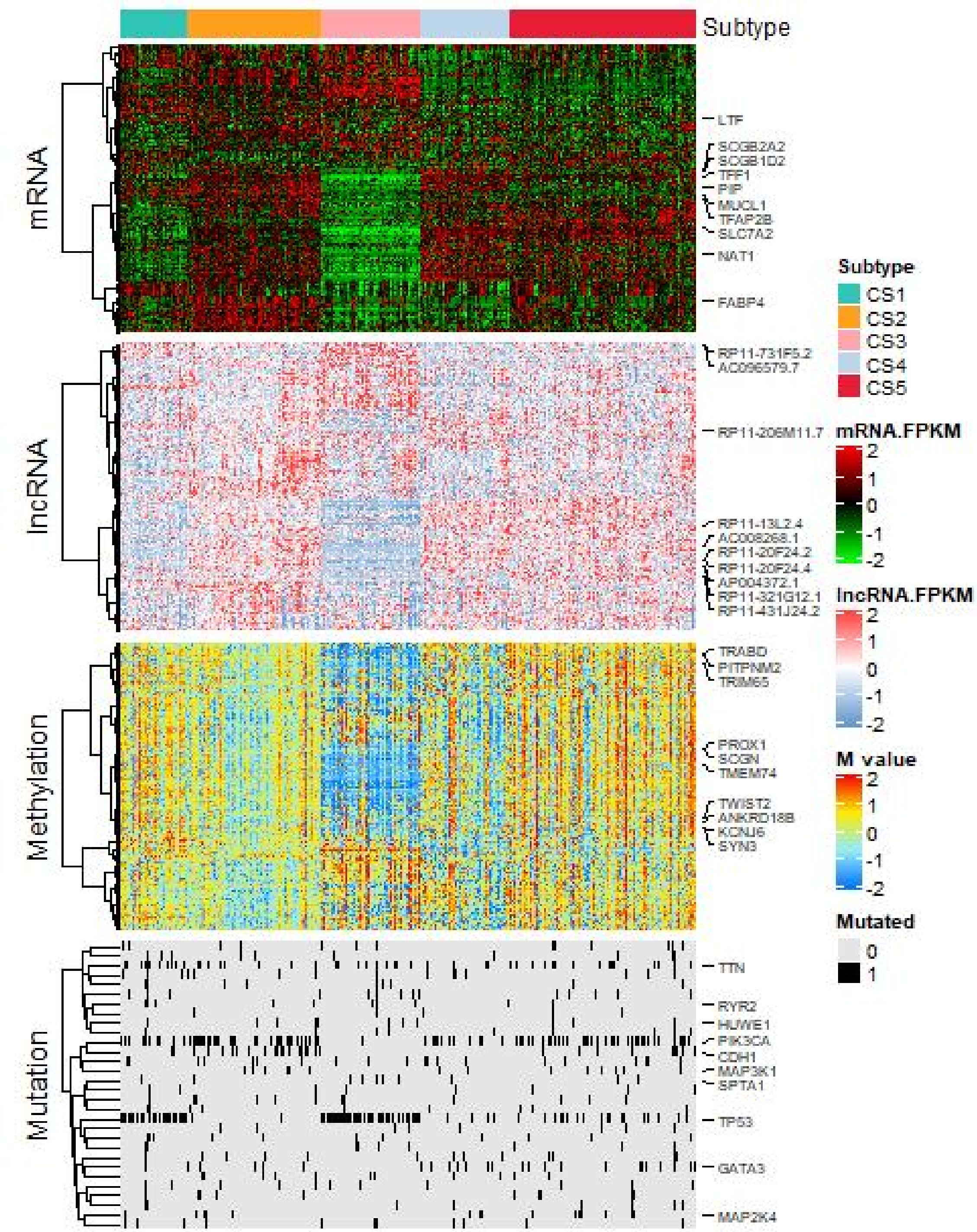

Figure 3: Comprehensive heatmap of multi-omics integrative clustering by iClusterBayes with annotation of potential drivers.

Of course, since there are a total of 10 results stored in the `moic.res.list`, you can also choose any one of them to create the heatmap. Here I select COCA which also returns sample dendrogram like below:

```
getMoHeatmap(data          = plotdata,
             row.title     = c("mRNA","lncRNA","Methylation","Mutation"),
             is.binary     = c(F,F,F,T),
             legend.name   = c("mRNA.FPKM","lncRNA.FPKM","M value","Mutated"),
             clust.res     = moic.res.list$COCA$clust.res, # cluster results
             clust.dend    = moic.res.list$COCA$clust.dend, # show sample dendrogram
             color         = col.list,
             width         = 10,
             height        = 5,
```

```
             fig.name       = "COMPREHENSIVE HEATMAP OF COCA")
```

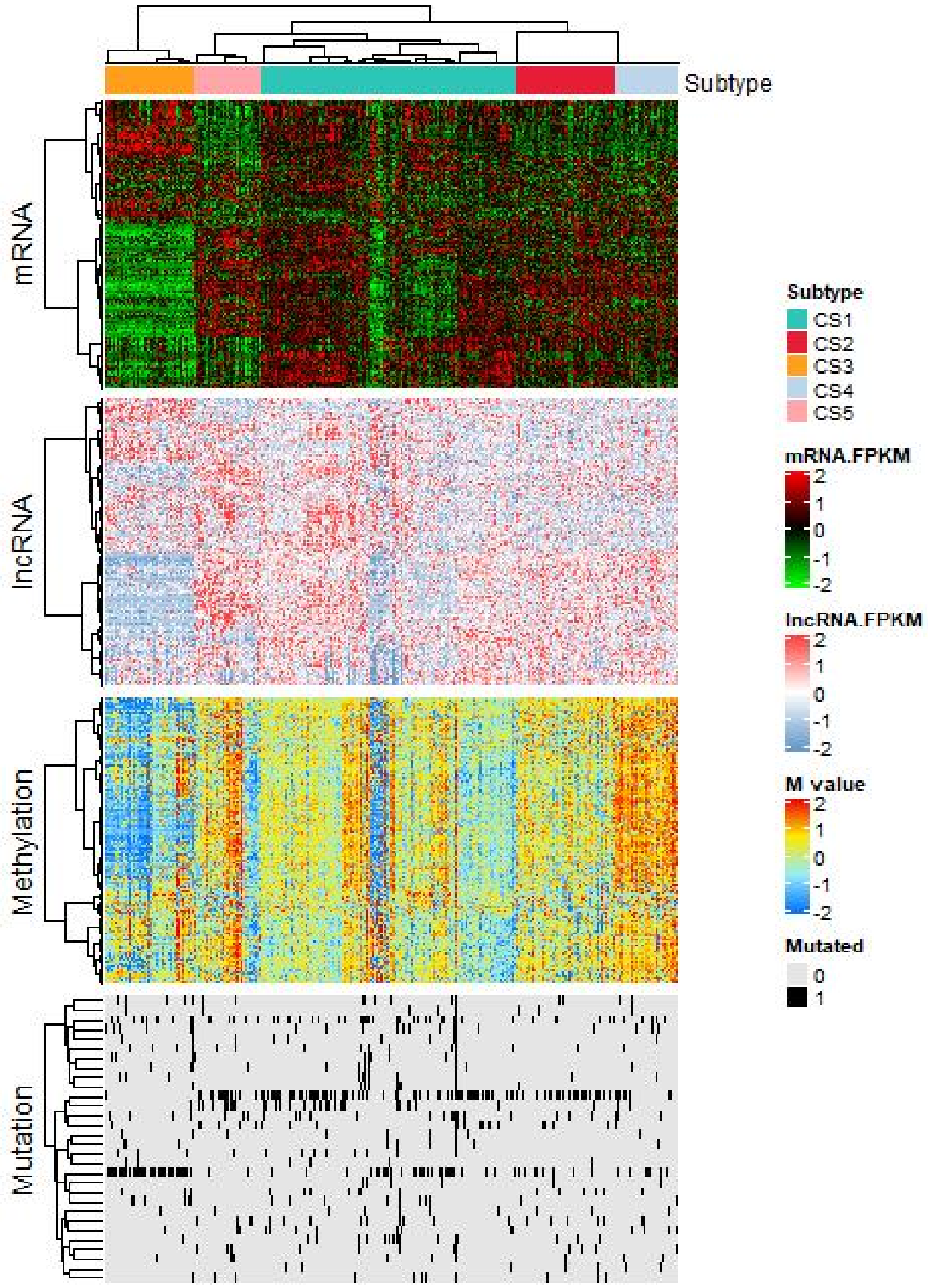

Figure 4: Comprehensive heatmap of multi-omics integrative clustering by COCA with dendrogram for samples.

Now go back to the consensus result of `cmoic.brca` that integrates 10 algorithms and this time samples' annotation is also provided to generate the heatmap. Since the core function of `getMoHeatmap()` is based on ComplexHeatmap R package[20], when creating annotations, we should always use `circlize::colorRamp2()` function to generate the color mapping function for continuous variables (*e.g.*, age in this example).

```
# extract PAM50, pathologic stage and age for sample annotation
annCol    <- surv.info[,c("PAM50", "pstage", "age"), drop = FALSE]
```

```r
# generate corresponding colors for sample annotation
annColors <- list(age    =
                    circlize::colorRamp2(breaks
                                         = c(min(annCol$age),
                                             median(annCol$age),
                                             max(annCol$age)),
                                         colors
                                         = c("#0000AA", "#555555", "#AAAA00")),
                  PAM50  = c("Basal" = "blue",
                             "Her2"   = "red",
                             "LumA"   = "yellow",
                             "LumB"   = "green",
                             "Normal" = "black"),
                  pstage = c("T1"    = "green",
                              "T2"    = "blue",
                              "T3"    = "red",
                              "T4"    = "yellow",
                              "TX"    = "black"))

# comprehensive heatmap
getMoHeatmap(data          = plotdata,
             row.title     = c("mRNA","lncRNA","Methylation","Mutation"),
             is.binary     = c(F,F,F,T),
             legend.name   = c("mRNA.FPKM","lncRNA.FPKM","M value","Mutated"),
             clust.res     = cmoic.brca$clust.res, # consensusMOIC results
             clust.dend    = NULL,
             show.rownames = c(F,F,F,F),
             show.colnames = FALSE,
             show.row.dend = c(F,F,F,F), # show no dendrogram for features
             annRow        = NULL, # no selected features
             color         = col.list,
             annCol        = annCol, # annotation for samples
             annColors     = annColors, # annotation color
             width         = 10,
             height        = 5,
             fig.name      = "COMPREHENSIVE HEATMAP OF CONSENSUSMOIC")
```

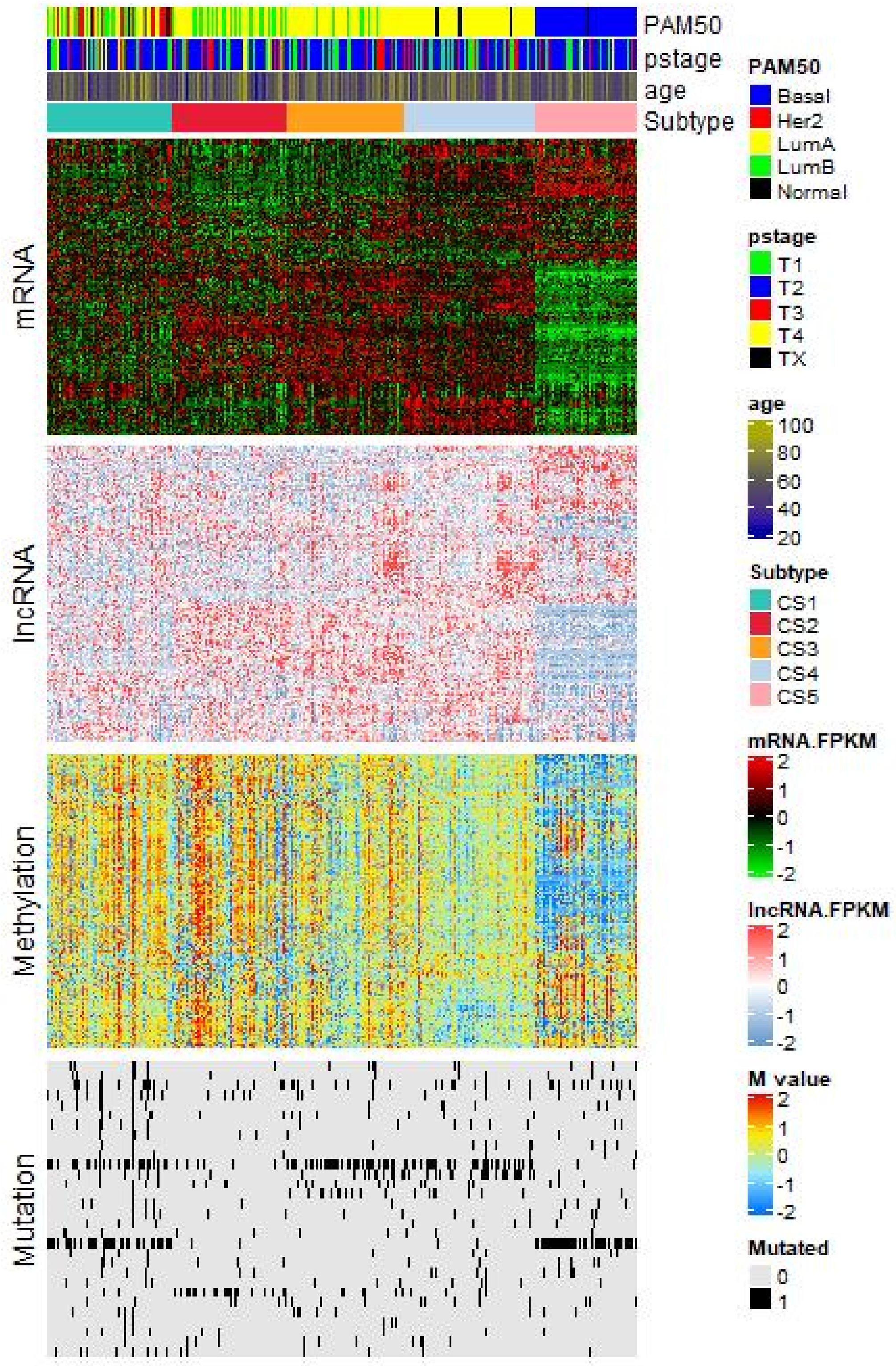

Figure 5: Comprehensive heatmap based on consensus across 10 algorithms with clinicopathological annotation.

###### 4.2.2 COMP Module

After identification of cancer subtypes, it is essential to further characterize each subtype by discovering their difference from multiple aspects. To this end, MOVICS provides commonly used downstream analyses in cancer subtyping researches for easily cohesion with results derived from **GET Module**.

###### 4.2.2.1 compare survival outcome

MOVICS provides function of `compSurv()` which not only calculates the overall nominal *P* value by log-rank test, but also performs pairwise comparison and derives adjusted *P* values if more than two subtypes are identified. These information will be all printed in the Kaplan-Meier Curve which is convenient for researchers to refer. Except for clustering results (*e.g.*, `cmoic.brca` in this example), you must additionally provide `surv.info` argument which should be a data.frame (must has row names of samples) that stores a `futime` column for survival time (**unit of day**) and another `fustat` column for survival outcome (0 = censor; 1 = event).

```
# survival comparison
surv.brca <-
  compSurv(moic.res         = cmoic.brca,
           surv.info        = surv.info,
           convt.time       = "m", # convert day unit to month
           surv.median.line = "h", # draw horizontal line at median survival
           fig.name         = "KAPLAN-MEIER CURVE OF CONSENSUSMOIC")
#> --a total of 643 samples are identified.
#> --removed missing values.
#> --leaving 642 observations.
#> --cut survival curve up to 10 years.
```

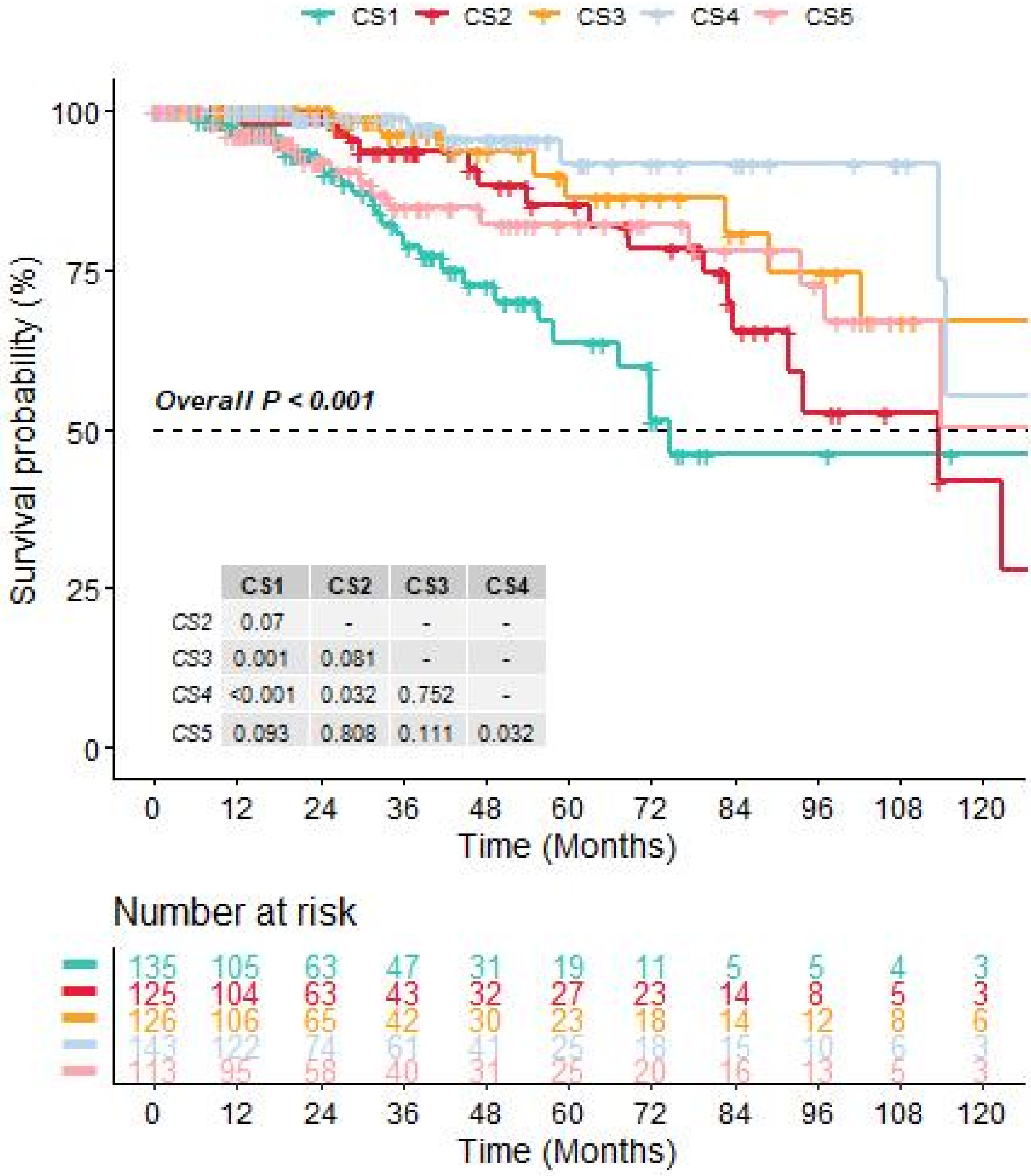

| | CS1 | CS2 | CS3 | CS4 |
|---|---|---|---|---|
| CS2 | 0.07 | - | - | - |
| CS3 | 0.001 | 0.081 | - | - |
| CS4 | <0.001 | 0.032 | 0.752 | - |
| CS5 | 0.093 | 0.808 | 0.111 | 0.032 |

| Time (Months) | 0 | 12 | 24 | 36 | 48 | 60 | 72 | 84 | 96 | 108 | 120 |
|---|---|---|---|---|---|---|---|---|---|---|---|
| CS1 | 135 | 105 | 63 | 47 | 31 | 19 | 11 | 5 | 5 | 4 | 3 |
| CS2 | 125 | 104 | 63 | 43 | 32 | 27 | 23 | 14 | 8 | 5 | 3 |
| CS3 | 126 | 106 | 65 | 42 | 30 | 23 | 18 | 14 | 12 | 8 | 6 |
| CS4 | 143 | 122 | 74 | 61 | 41 | 25 | 18 | 15 | 10 | 6 | 3 |
| CS5 | 113 | 95 | 58 | 40 | 31 | 25 | 20 | 16 | 13 | 5 | 3 |

Figure 6: Kaplan-Meier survival curve of 5 identified subtypes of breast cancer in TCGA-BRCA cohort.

```
print(surv.brca)
#> $fitd
#> Call:
#> survdiff(formula = Surv(futime, fustat) ~ Subtype, data = mosurv.res,
#>     na.action = na.exclude)
#>
#>                  N Observed Expected (O-E)^2/E (O-E)^2/V
#> Subtype=CS1 135       27     12.7    16.078    19.640
#> Subtype=CS2 125       17     14.6     0.396     0.496
#> Subtype=CS3 126        8     15.9     3.909     5.033
#> Subtype=CS4 143        7     17.2     6.089     7.982
#> Subtype=CS5 113       16     14.6     0.140     0.176
#>
#>  Chisq= 27.1  on 4 degrees of freedom, p= 2e-05
```

```
#>
#> $fit
#> Call: survfit(formula = Surv(futime, fustat) ~ Subtype, data = mosurv.res,
#>     na.action = na.exclude, error = "greenwood", type = "kaplan-meier",
#>     conf.type = "plain")
#>
#>       n events median 0.95LCL 0.95UCL
#> CS1 135     27   74.5    57.7      NA
#> CS2 125     17  113.5    83.6     144
#> CS3 126      8     NA   102.5      NA
#> CS4 143      7  216.2   113.5      NA
#> CS5 113     16     NA    97.2      NA
#>
#> $overall.p
#> [1] 1.918496e-05
#>
#> $pairwise.p
#>
#>  Pairwise comparisons using Log-Rank test
#>
#> data:  mosurv.res and Subtype
#>
#>     CS1     CS2     CS3     CS4
#> CS2 0.06963 -       -       -
#> CS3 0.00142 0.08079 -       -
#> CS4 0.00018 0.03205 0.75191 -
#> CS5 0.09345 0.80850 0.11121 0.03205
#>
#> P value adjustment method: BH
```

**4.2.2.2 compare clinical features** We then compare the clinical features among different subtypes. MOVICS provides function of `compClinvar()` which enables summarizing both continuous and categorical variables and performing proper statistical tests. This function can give a table in .docx format that is easy to use in medical research papers.

```
clin.brca <-
  compClinvar(moic.res      = cmoic.brca,
              var2comp      = surv.info, # data.frame needs to summarize
              strata        = "Subtype", # stratifying variable
              factorVars    = c("PAM50","pstage","fustat"), # categorical variables
              nonnormalVars = "futime", # feature(s) to use nonparametric test
              exactVars     = "pstage", # feature(s) to use exact test
              doWord        = TRUE, # generate .docx file in local path
              tab.name      = "SUMMARIZATION OF CLINICAL FEATURES")
#> --all samples matched.
#> Warning in jstable::CreateTableOne2(vars = setdiff(colnames(dat), strata), : NAs
#> introduced by coercion
```

```
print(clin.brca$compTab)
```

**4.2.2.3 compare mutational frequency** Subype-specific mutation might be promising as therapeutic target. Hence, we then compare the mutational frequency among different clusters. MOVICS provides function of `compMut()` which deals with binary mutational data (0 = wild; 1 = mutated). This function applies independent testing (*i.e.*, Fisher' s exact test or $\chi^2$ test) for each mutation, and generates a table with statistical results (.docx file also if specified), and creates an OncoPrint using those differentially mutated genes.

Table 1: Comparison of clinical features among 5 identified subtype of breast cancer in TCGA-BRCA cohort.

| | level | CS1 | CS2 | CS3 | CS4 | CS5 | p | test | sig |
|---|---|---|---|---|---|---|---|---|---|
| n | | 135 | 126 | 126 | 143 | 113 | | | |
| fustat (%) | 0 | 108 (80.0) | 108 (85.7) | 118 (93.7) | 136 (95.1) | 97 (85.8) | <0.001 | | ** |
| | 1 | 27 (20.0) | 18 (14.3) | 8 ( 6.3) | 7 ( 4.9) | 16 (14.2) | | | |
| futime (median [IQR]) | | 675.00 [404.00, 1317.00] | 746.00 [435.00, 1528.00] | 760.50 [444.75, 1316.25] | 837.00 [465.00, 1546.00] | 742.00 [470.00, 1605.00] | 0.602 | nonnorm | |
| PAM50 (%) | Basal | 2 ( 1.5) | 0 ( 0.0) | 0 ( 0.0) | 0 ( 0.0) | 109 (96.5) | <0.001 | | ** |
| | Her2 | 38 (28.1) | 0 ( 0.0) | 0 ( 0.0) | 0 ( 0.0) | 0 ( 0.0) | | | |
| | LumA | 39 (28.9) | 72 (57.1) | 106 (84.1) | 127 (88.8) | 0 ( 0.0) | | | |
| | LumB | 50 (37.0) | 54 (42.9) | 20 (15.9) | 0 ( 0.0) | 0 ( 0.0) | | | |
| | Normal | 6 ( 4.4) | 0 ( 0.0) | 0 ( 0.0) | 16 (11.2) | 4 ( 3.5) | | | |
| pstage (%) | T1 | 37 (27.4) | 33 (26.2) | 37 (29.4) | 40 (28.0) | 25 (22.1) | NA | exact | |
| | T2 | 80 (59.3) | 74 (58.7) | 72 (57.1) | 71 (49.7) | 70 (61.9) | | | |
| | T3 | 10 ( 7.4) | 15 (11.9) | 14 (11.1) | 31 (21.7) | 14 (12.4) | | | |
| | T4 | 8 ( 5.9) | 4 ( 3.2) | 3 ( 2.4) | 1 ( 0.7) | 3 ( 2.7) | | | |
| | TX | 0 ( 0.0) | 0 ( 0.0) | 0 ( 0.0) | 0 ( 0.0) | 1 ( 0.9) | | | |
| age | | 58.53 (13.41) | 59.63 (14.10) | 59.16 (12.90) | 57.53 (12.26) | 55.67 (12.11) | 0.127 | | |

```
# mutational frequency comparison
mut.brca <-
  compMut(moic.res     = cmoic.brca,
          mut.matrix   = brca.tcga$mut.status, # binary somatic mutation matrix
          doWord       = TRUE, # generate table in .docx format
          doPlot       = TRUE, # draw OncoPrint
          freq.cutoff  = 0.05, # keep genes that mutated in at least 5% of samples
          p.adj.cutoff = 0.05, # keep genes with adjusted pvalue<0.05 in OncoPrint
          innerclust   = TRUE, # perform clustering within each subtype
          annCol       = annCol, # same annotation for heatmap
          annColors    = annColors, # same annotation color for heatmap
          width        = 6,
          height       = 2,
          fig.name     = "ONCOPRINT FOR SIGNIFICANT MUTATIONS",
          tab.name     = "INDEPENDENT TEST BETWEEN SUBTYPE AND MUTATION")
#> --all samples matched.
```

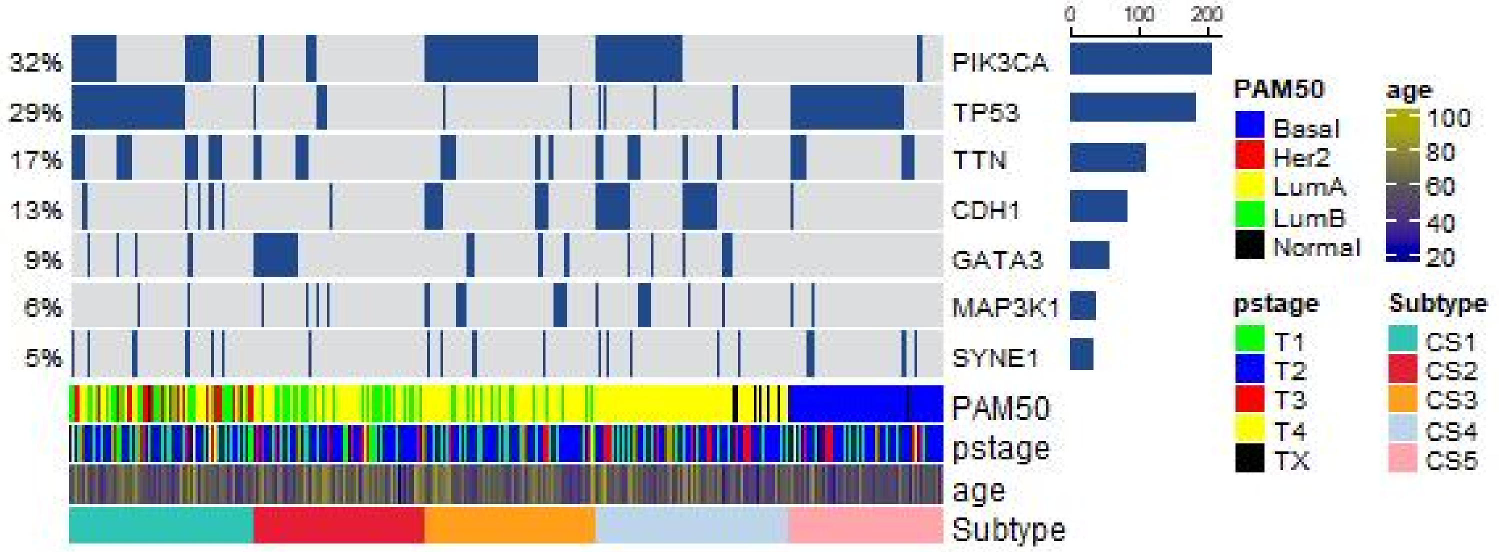

Figure 7: Mutational OncoPrint of 5 identified subtypes of breast cancer in TCGA-BRCA cohort.

```
print(mut.brca)
```

Table 2: Comparison of mutational frequency among 5 identified subtype of breast cancer in TCGA-BRCA cohort.

| Gene (Mutated) | TMB | CS1 | CS2 | CS3 | CS4 | CS5 | pvalue | padj |
|---|---|---|---|---|---|---|---|---|
| PIK3CA | 208 (32%) | 51 (37.8%) | 9 ( 7.1%) | 82 (65.1%) | 64 (44.8%) | 2 ( 1.8%) | 9.08e-39 | 4.09e-38 |
| TP53 | 186 (29%) | 84 (62.2%) | 9 ( 7.1%) | 2 ( 1.6%) | 8 ( 5.6%) | 83 (73.5%) | 1.91e-67 | 1.72e-66 |
| TTN | 111 (17%) | 40 (29.6%) | 12 ( 9.5%) | 17 (13.5%) | 22 (15.4%) | 20 (17.7%) | 4.75e-04 | 7.13e-04 |
| CDH1 | 83 (13%) | 9 ( 6.7%) | 3 ( 2.4%) | 21 (16.7%) | 49 (34.3%) | 1 ( 0.9%) | 6.84e-19 | 2.05e-18 |
| GATA3 | 58 ( 9%) | 5 ( 3.7%) | 31 (24.6%) | 11 ( 8.7%) | 11 ( 7.7%) | 0 ( 0.0%) | 1.08e-10 | 2.43e-10 |
| MLL3 | 49 ( 8%) | 11 (8.1%) | 12 (9.5%) | 10 (7.9%) | 11 (7.7%) | 5 (4.4%) | 6.55e-01 | 6.55e-01 |
| MUC16 | 48 ( 8%) | 16 (11.9%) | 8 ( 6.3%) | 8 ( 6.3%) | 8 ( 5.6%) | 8 ( 7.1%) | 3.48e-01 | 3.91e-01 |
| MAP3K1 | 38 ( 6%) | 2 ( 1.5%) | 5 ( 4.0%) | 17 (13.5%) | 12 ( 8.4%) | 2 ( 1.8%) | 1.15e-04 | 2.07e-04 |
| SYNE1 | 33 ( 5%) | 9 (6.7%) | 1 (0.8%) | 5 (4.0%) | 8 (5.6%) | 10 (8.8%) | 3.24e-02 | 4.17e-02 |

**4.2.2.4 compare total mutation burden** Needless to say, immunotherapy is becoming a pillar of modern cancer treatment. Recent analyses have linked the tumoral genomic landscape with antitumor immunity. In particular, it has been proposed that overall mutational load might drive T-cell responses[21], whereas tumor aneuploidy correlates with markers of immune evasion and reduced response to immunotherapy[22]. To quantify these genomic alterations that may affect immunotherapy, MOVICS provides two functions to calculate total mutation burden (TMB) and fraction genome altered (FGA). To be specific, Total Mutations Burden (TMB) refers to the number of mutations that are found in the tumor genome[23,24], while FGA is the percentage of genome that has been affected by copy number gains or losses[25,26]. Both attributes are useful for genetic researchers as they provide them with more in-depth information on the genomic make-up of the tumors. Let me show you how to use these two functions by starting with `compTMB()`. First of all, the input maf data for this function must have the following 10 columns at least:

```
head(maf)
#>   Tumor_Sample_Barcode Hugo_Symbol Chromosome Start_Position End_Position
#> 1         BRCA-A1XY-01A       USP24       chr1       55159655     55159655
#> 2         BRCA-A1XY-01A      ERICH3       chr1       74571494     74571494
#> 3         BRCA-A1XY-01A      KIF26B       chr1      245419680    245419680
#> 4         BRCA-A1XY-01A       USP34       chr2       61189055     61189055
#> 5         BRCA-A1XY-01A      ANTXR1       chr2       69245305     69245305
#> 6         BRCA-A1XY-01A       SCN9A       chr2      166199365    166199365
#>   Variant_Classification Variant_Type Reference_Allele Tumor_Seq_Allele1
```

```
#> 1      Missense_Mutation          SNP                T                 T
#> 2      Missense_Mutation          SNP                C                 C
#> 3                 Silent          SNP                G                 G
#> 4                 Silent          SNP                G                 G
#> 5                 Silent          SNP                G                 G
#> 6                 Silent          SNP                G                 G
#>   Tumor_Seq_Allele2
#> 1                 C
#> 2                 T
#> 3                 T
#> 4                 C
#> 5                 A
#> 6                 A
```

By default, `compTMB()` considers nonsynonymous variants only when counting somatic mutation frequency, including Frame_Shift_Del, Frame_Shift_Ins, Splice_Site, Translation_Start_Site, Nonsense_Mutation, Nonstop_Mutation, In_Frame_Del, In_Frame_Ins, and Missense_Mutation. In addition to calculating TMB, this function also classifies Single Nucleotide Variants into Transitions and Transversions (TiTv), and depicts the distribution of both TMB and TiTv.

```
# compare TMB
tmb.brca <-
  compTMB(moic.res     = cmoic.brca,
          maf          = maf,
          rmDup        = TRUE, # remove duplicated variants per sample
          rmFLAGS      = FALSE, # keep FLAGS mutations
          exome.size   = 38, # estimated exome size
          test.method  = "nonparametric", # statistical testing method
          fig.name     = "DISTRIBUTION OF TMB AND TITV")
#> --67 samples mismatched from current subtypes.
#> -Validating
#> -Silent variants: 24329
#> -Summarizing
#> --Possible FLAGS among top ten genes:
#>   TTN
#>   MUC16
#> -Processing clinical data
#> --Missing clinical data
#> -Finished in 3.420s elapsed (3.420s cpu)
```

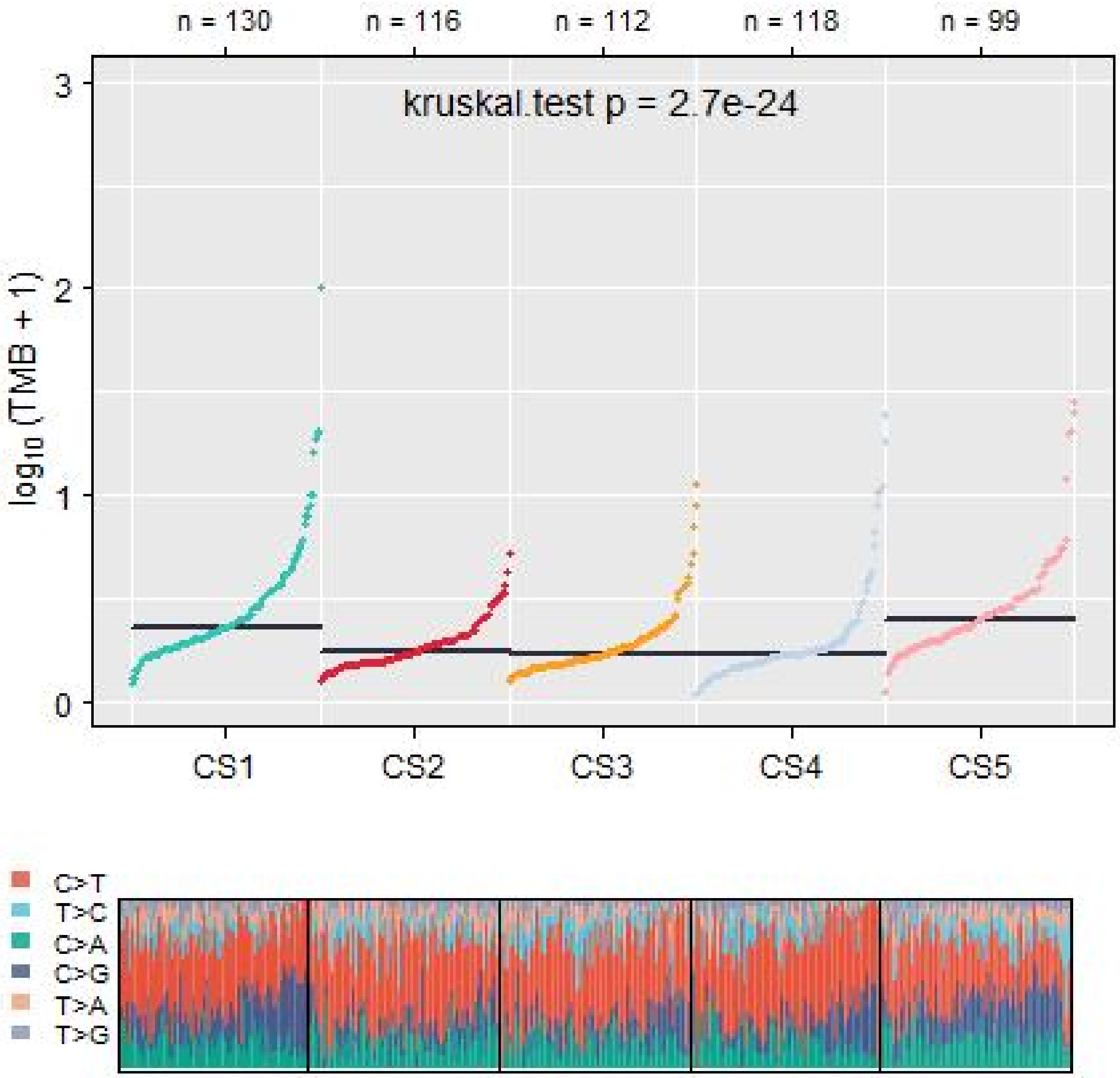

Figure 8: Comparison of TMB and TiTv among 5 identified subtypes of breast cancer in TCGA-BRCA cohort.

```
head(tmb.brca$TMB.dat)
```

Table 3: Demo of comparison of TMB among 5 identified subtype of breast cancer in TCGA-BRCA cohort.

| | samID | variants | TMB | log10TMB | Subtype |
|---|---|---|---|---|---|
| 570 | BRCA-A1EW-01A | 9 | 0.2368421 | 0.0923143 | CS1 |
| 558 | BRCA-A1IO-01A | 11 | 0.2894737 | 0.1104125 | CS1 |
| 560 | BRCA-A0C3-01A | 11 | 0.2894737 | 0.1104125 | CS1 |
| 525 | BRCA-A5RY-01A | 16 | 0.4210526 | 0.1526102 | CS1 |
| 528 | BRCA-A1EX-01A | 16 | 0.4210526 | 0.1526102 | CS1 |
| 506 | BRCA-A1FE-01A | 18 | 0.4736842 | 0.1684044 | CS1 |

**4.2.2.5 compare fraction genome altered** Next, `compFGA()` calculates not only FGA but also computes specific gain (FGG) or loss (FGL) per sample within each subtype. To get this function worked, an eligible input of segmented copy number should be prepared with exactly the same column name like below:

```r
# change column names of segment data
colnames(segment) <- c("sample","chrom","start","end","value")
```

Let's see how the input looks like:

```r
head(segment)
#>          sample chrom     start       end   value
#> 1 BRCA-A090-01A     1   3301765  54730235 -0.1271
#> 2 BRCA-A090-01A     1  54730247  57443819 -0.0899
#> 3 BRCA-A090-01A     1  57448465  57448876 -1.1956
#> 4 BRCA-A090-01A     1  57448951  64426102 -0.1009
#> 5 BRCA-A090-01A     1  64426648 106657734 -0.1252
#> 6 BRCA-A090-01A     1 106657854 106835667  0.1371
```

Notably, if your CNA calling procedure did not provide a segmented copy number as value column but the original copy number, argument of `iscopynumber` must be switched to `TRUE` instead.

```r
# compare FGA, FGG, and FGL
fga.brca <-
  compFGA(moic.res      = cmoic.brca,
          segment       = segment,
          iscopynumber = FALSE, # this is a segmented copy number file
          cnathreshold = 0.2, # threshold to determine CNA gain or loss
          test.method  = "nonparametric", # statistical testing method
          fig.name     = "BARPLOT OF FGA")
#> --2 samples mismatched from current subtypes.
#> 5% 10% 15% 21% 26% 31% 36% 41% 46% 51% 57% 62% 67% 72% 77% 82% 88% 93% 98%
```

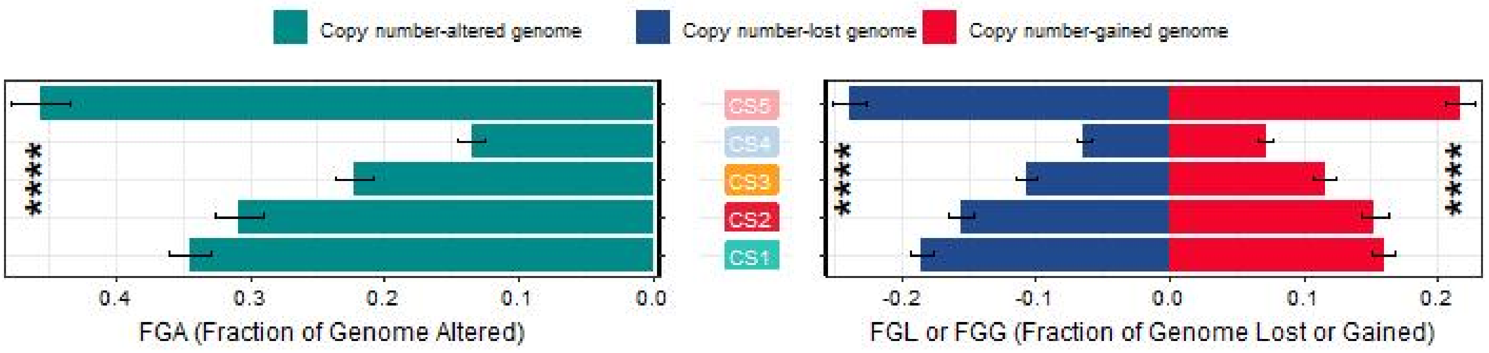

Figure 9: Barplot of fraction genome altered among 5 identified subtypes of breast cancer in TCGA-BRCA cohort.

```r
head(fga.brca$summary)
```

Table 4: Demo of comparison of fraction genome altered among 5 identified subtype of breast cancer in TCGA-BRCA cohort.

| samID | FGA | FGG | FGL | Subtype |
|---|---|---|---|---|
| BRCA-A03L-01A | 0.6217991 | 0.3086727 | 0.3131264 | CS1 |
| BRCA-A04R-01A | 0.2531019 | 0.0913201 | 0.1617818 | CS2 |
| BRCA-A075-01A | 0.7007067 | 0.4144424 | 0.2862643 | CS1 |
| BRCA-A08O-01A | 0.6501287 | 0.4564814 | 0.1936473 | CS3 |
| BRCA-A0A6-01A | 0.1468893 | 0.0635649 | 0.0833244 | CS4 |
| BRCA-A0AD-01A | 0.1722214 | 0.0386452 | 0.1335762 | CS2 |

**4.2.2.6 compare drug sensitivity** Predicting response to medication is particularly important for drugs with a narrow therapeutic index, for example chemotherapeutic agents, because response is highly variable and side effects are potentially lethal. Therefore, Paul Geeleher et al. (2014)[27] used baseline gene expression and *in vitro* drug sensitivity derived from cell lines, coupled with *in vivo* baseline tumor gene expression, to predict patients' response to drugs. Paul developped an R package pRRophetic for prediction of clinical chemotherapeutic response from tumor gene expression levels[28], and now this function has been involved in MOVICS to examine difference of drug sensitivity among different subtypes. Here we estimate the $IC_{50}$ of *Cisplatin* and *Paclitaxel* for 5 identified subtypes of breast cancer in TCGA cohort.

```
# drug sensitivity comparison
drug.brca <-
  compDrugsen(moic.res    = cmoic.brca,
              norm.expr   = fpkm[,cmoic.brca$clust.res$samID],
              drugs       = c("Cisplatin", "Paclitaxel"), # a vector of drug name
              tissueType  = "breast", # choose specific tissue type
              test.method = "nonparametric", # statistical testing method
              prefix      = "BOXVIOLIN OF ESTIMATED IC50")
#> --all samples matched.
#> --log2 transformation done for expression data.
#> Cisplatin done...
#> Paclitaxel done...
```

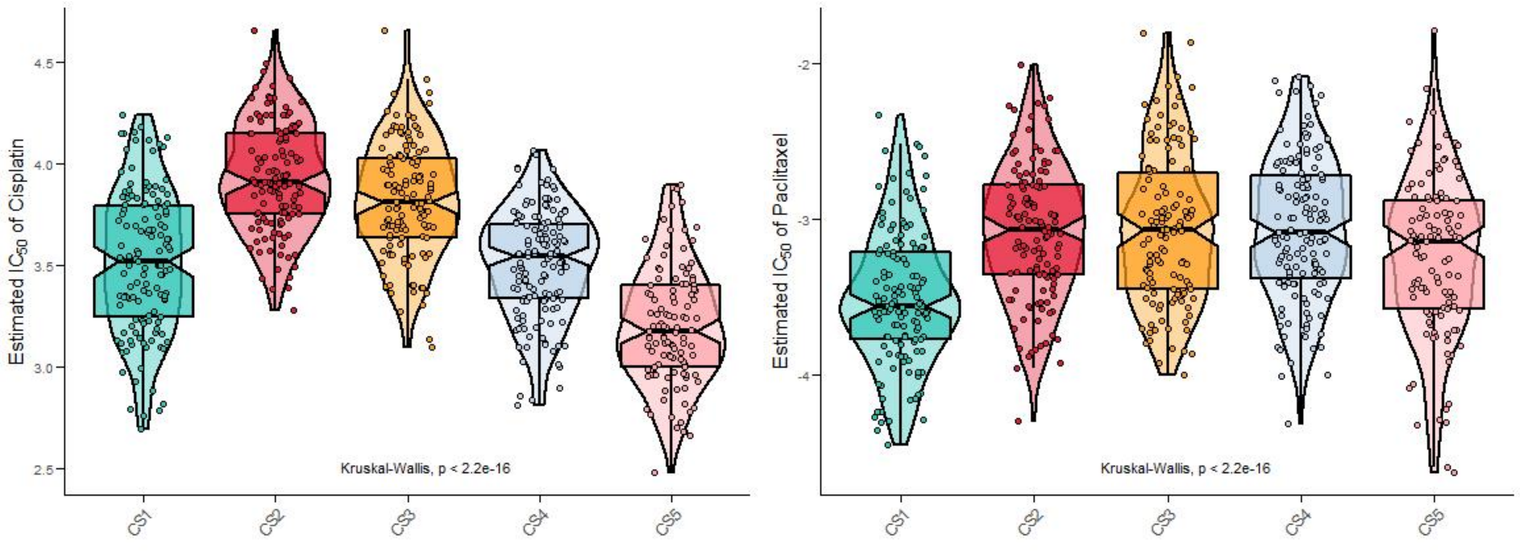

Figure 10: Boxviolins for estimated IC50 of Cisplatin and Paclitaxel among 5 identified subtypes of breast cancer in TCGA-BRCA cohort.

```
head(drug.brca$Cisplatin)
```

Table 5: Demo of estimated IC50 for Cisplatin among 5 identified subtype of breast cancer in TCGA-BRCA cohort.

| | Est.IC50 | Subtype |
|---|---|---|
| BRCA-A03L-01A | 3.802060 | CS1 |
| BRCA-A075-01A | 3.834646 | CS1 |
| BRCA-A0AW-01A | 3.126744 | CS1 |
| BRCA-A0BC-01A | 3.409402 | CS1 |
| BRCA-A0BF-01A | 3.358053 | CS1 |
| BRCA-A0BT-01A | 4.239210 | CS1 |

**4.2.2.7 compare agreement with other subtypes** At present, many cancers have traditional classifications, and evaluating the consistency of new subtypes with previous classifications is critical to reflect the robustness of clustering analysis and to determine potential but novel subtypes. To measure the agreement (similarity) between the current subtypes and other pre-existed classifications, MOVICS provides function of `compAgree()` to calculate four statistics: Rand Index (RI)[29], Adjusted Mutual Information (AMI)[30], Jaccard Index (JI)[31], and Fowlkes-Mallows (FM)[32]; all these measurements range from 0 to 1 and the larger the value is, the more similar the two appraises are. This function can also generate an alluvial diagram to visualize the agreement of two appraises with the current subtypes as reference.

```
# customize the factor level for pstage
surv.info$pstage <- factor(surv.info$pstage, levels = c("TX","T1","T2","T3","T4"))

# agreement comparison (support up to 6 classifications include current subtype)
agree.brca <-
  compAgree(moic.res  = cmoic.brca,
            subt2comp = surv.info[,c("PAM50","pstage")],
            doPlot    = TRUE,
            box.width = 0.2,
            fig.name  = "AGREEMENT OF CONSENSUSMOIC WITH PAM50 AND PSTAGE")
#> --all samples matched.
```

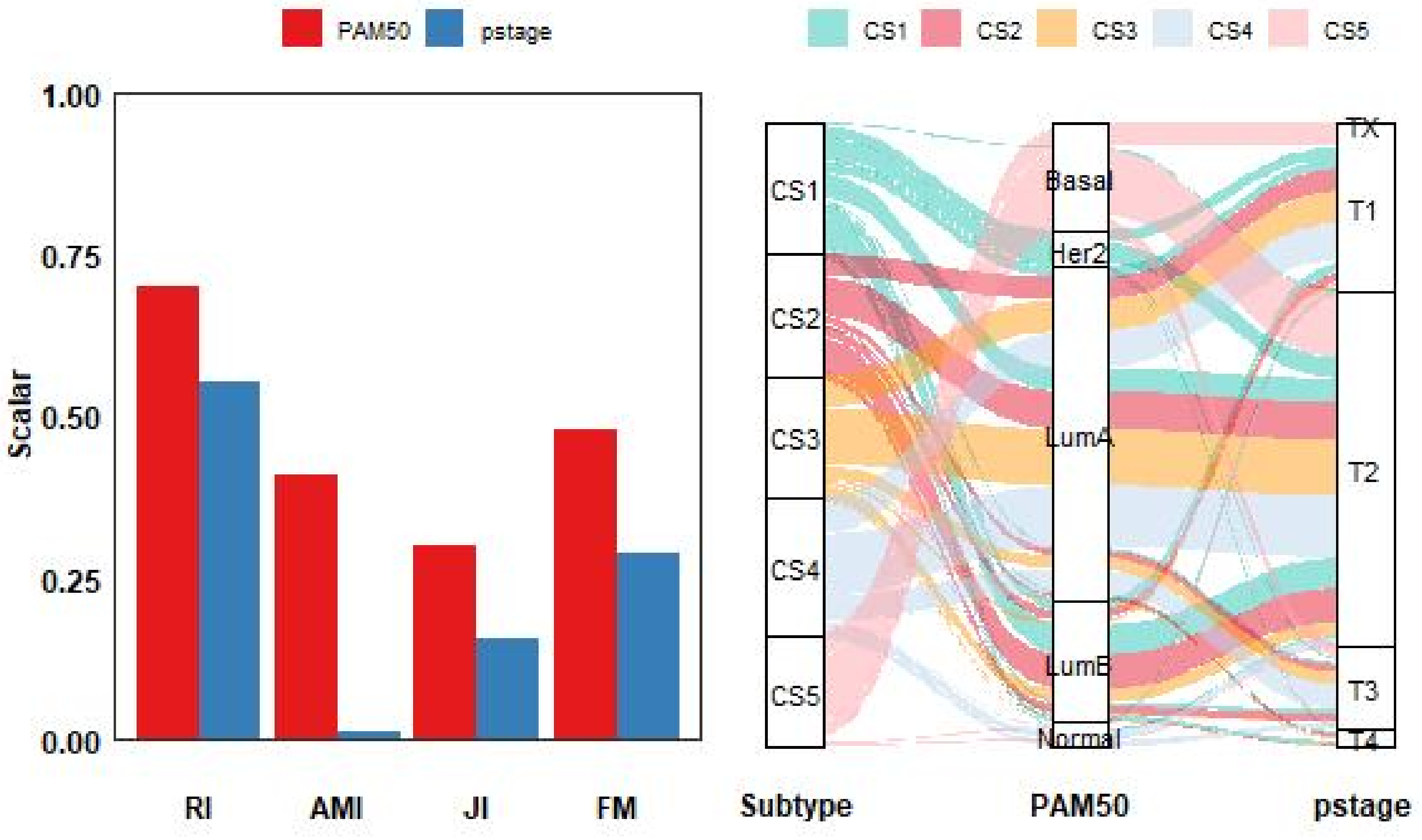

Figure 11: Agreement of 5 identified subtypes of breast cancer with PAM50 classification and pathological stage in TCGA-BRCA cohort.

```
print(agree.brca)
```

Table 6: Agreement of 5 identified subtypes with PAM50 classification and pathological stage in TCGA-BRCA cohort.

| current.subtype | other.subtype | RI | AMI | JI | FM |
|---|---|---|---|---|---|
| Subtype | PAM50 | 0.6988852 | 0.4097097 | 0.2984502 | 0.4792001 |
| Subtype | pstage | 0.5513050 | 0.0121390 | 0.1558165 | 0.2877073 |

###### 4.2.3 RUN Module

In **RUN Module**, MOVICS aims to characterize different subtypes by identifying their potential predictive biomarkers and functional pathways. Identifying and applying molecular biomarkers to predict subtype with efficiency is particularly important for disease management and treatment, thus improving clinical outcome. In this context, MOVICS searches for subtype-specific biomarkers by starting with differential expression analysis (DEA).

**4.2.3.1 run differential expression analysis** MOVICS provides `runDEA()` function which embeds three state-of-the-art DEA approches to identify differentially expressed genes (DEGs), including edgeR[33,34] and DESeq2[35] for RNA-Seq count data and limma[36] for microarray profile or normalized expression data. Since `runDEA()` checks the data scale automatically when choosing limma algorithm, it is recommended to provide a microarray expression profile or normalized expression data (*e.g.*, RSEM, FPKM, TPM) without z-score or log2 transformation.

```
# run DEA with edgeR
runDEA(dea.method = "edger",
       expr       = count, # raw count data
       moic.res   = cmoic.brca,
       prefix     = "TCGA-BRCA") # prefix of figure name
#> --all samples matched.
#> --you choose edger and please make sure an RNA-Seq count data was provided.
#> edger of CS1_vs_Others done...
#> edger of CS2_vs_Others done...
#> edger of CS3_vs_Others done...
#> edger of CS4_vs_Others done...
#> edger of CS5_vs_Others done...

# run DEA with DESeq2
runDEA(dea.method = "deseq2",
       expr       = count,
       moic.res   = cmoic.brca,
       prefix     = "TCGA-BRCA")
#> --all samples matched.
#> --you choose deseq2 and please make sure an RNA-Seq count data was provided.
#> deseq2 of CS1_vs_Others done...
#> deseq2 of CS2_vs_Others done...
#> deseq2 of CS3_vs_Others done...
#> deseq2 of CS4_vs_Others done...
#> deseq2 of CS5_vs_Others done...

# run DEA with limma
runDEA(dea.method = "limma",
       expr       = fpkm, # normalized expression data
       moic.res   = cmoic.brca,
       prefix     = "TCGA-BRCA")
#> --all samples matched.
#> --you choose limma and please make sure a microarray profile or a normalized expression data [FPKM or
#> --log2 transformation done for expression data.
#> limma of CS1_vs_Others done...
#> limma of CS2_vs_Others done...
```

```
#> limma of CS3_vs_Others done...
#> limma of CS4_vs_Others done...
#> limma of CS5_vs_Others done...
```

Each identified cancer subtype will be compared with the rest (Others) and the corresponding .txt file will be stored according to the argument of `res.path`. By default, these files will be saved under the current working directory.

###### 4.2.3.2 run biomarker identification procedure

In this procedure, the most differentially expressed genes sorted by log2FoldChange are chosen as the biomarkers for each subtype (200 biomarkers for each subtype by default). These biomarkers should pass the significance threshold (*e.g.*, nominal $P$ value < 0.05 and adjusted $P$ value < 0.05) and must not overlap with any biomarkers identified for other subtypes.

```
# choose edgeR result to identify subtype-specific up-regulated biomarkers
marker.up <-
  runMarker(moic.res      = cmoic.brca,
            dea.method    = "edger", # name of DEA method
            prefix        = "TCGA-BRCA", # MUST be the same of argument in runDEA()
            dat.path      = getwd(), # path of DEA files
            res.path      = getwd(), # path to save marker files
            p.cutoff      = 0.05, # p cutoff to identify significant DEGs
            p.adj.cutoff  = 0.05, # padj cutoff to identify significant DEGs
            dirct         = "up", # direction of dysregulation in expression
            n.marker      = 100, # number of biomarkers for each subtype
            doplot        = TRUE, # generate diagonal heatmap
            norm.expr     = fpkm, # use normalized expression as heatmap input
            annCol        = annCol, # sample annotation in heatmap
            annColors     = annColors, # colors for sample annotation
            show_rownames = FALSE, # show no rownames (biomarker name)
            fig.name      = "UPREGULATED BIOMARKER HEATMAP")
#> --all samples matched.
#> --log2 transformation done for expression data.
```

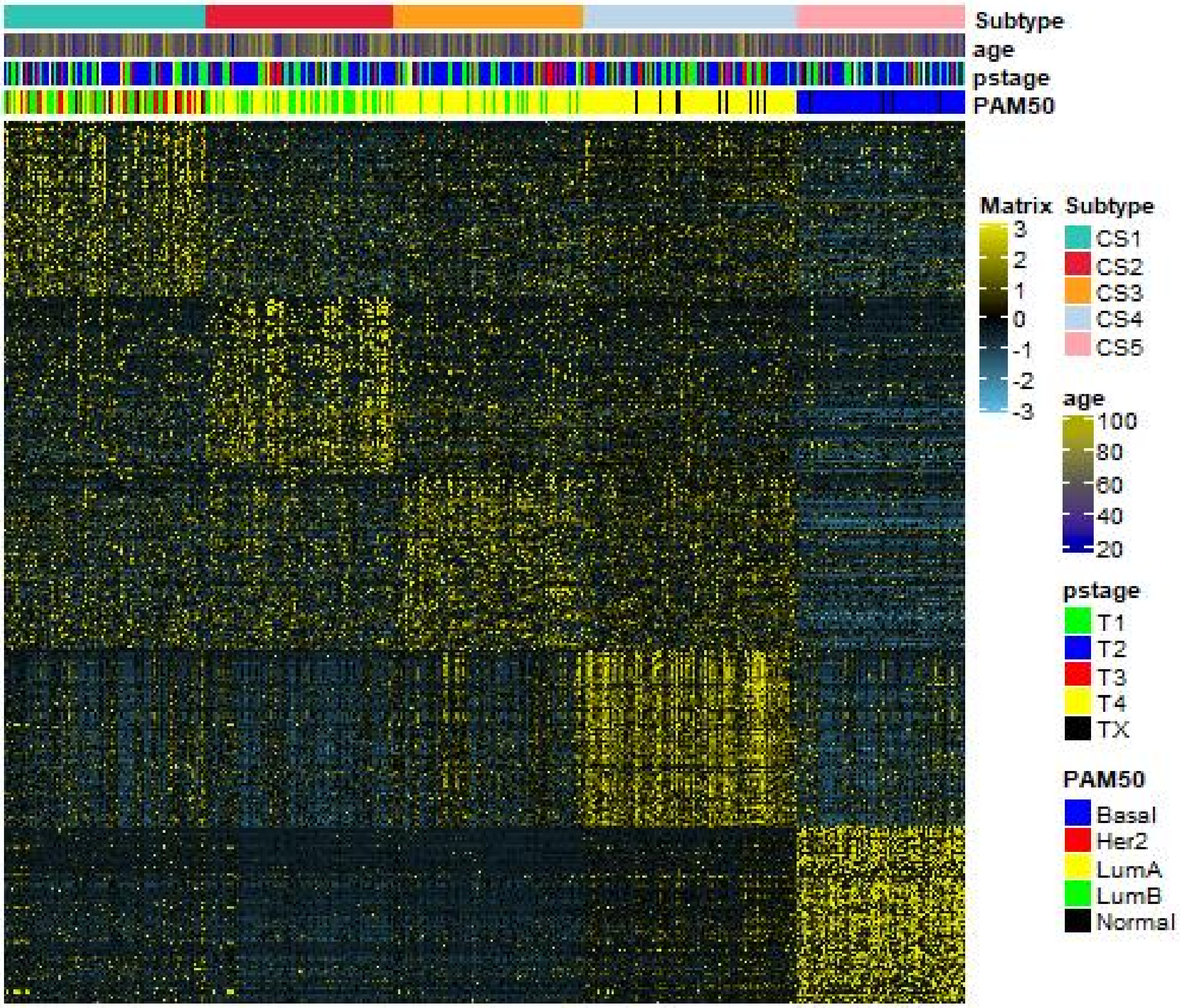

Figure 12: Heatmap of subtype-specific upregulated biomarkers using edgeR for 5 identified subtypes in TCGA-BRCA cohort.

```
# check the upregulated biomarkers
head(marker.up$templates)
```

Table 7: Demo of subtype-specific upregulated biomarkers for 5 identified subtypes of breast cancer in TCGA-BRCA cohort.

| probe | class | dirct |
|---|---|---|
| PNMT | CS1 | up |
| AKR1B15 | CS1 | up |
| DLK1 | CS1 | up |
| ACE2 | CS1 | up |
| CRISP3 | CS1 | up |
| ACSM1 | CS1 | up |

Then try results derived from limma (or DESeq2 if you want) to identify subtype-specific downregulated biomarkers.

```
# choose limma result to identify subtype-specific down-regulated biomarkers
marker.dn <-
  runMarker(moic.res       = cmoic.brca,
            dea.method     = "limma",
```

```
          prefix        = "TCGA-BRCA",
          dirct         = "down",
          n.marker      = 50, # switch to 50
          doplot        = TRUE,
          norm.expr     = fpkm,
          annCol        = annCol,
          annColors     = annColors,
          fig.name      = "DOWNREGULATED BIOMARKER HEATMAP")
#> --all samples matched.
#> --log2 transformation done for expression data.
```

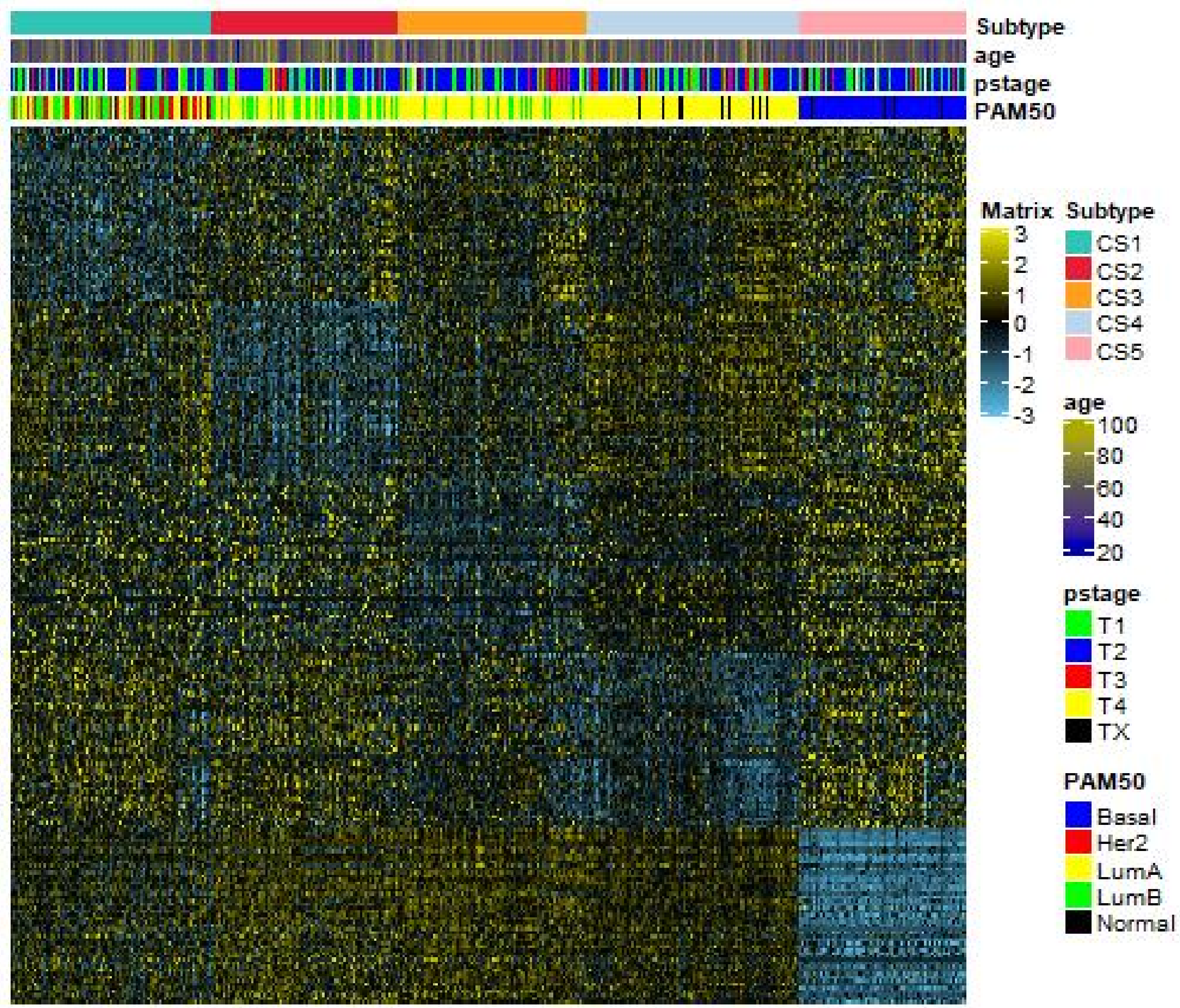

Figure 13: Heatmap of subtype-specific downregulated biomarkers using limma for 5 identified subtypes in TCGA-BRCA cohort.

**4.2.3.3 run gene set enrichment analysis** Similarly, GSEA[37] is run for each subtype based on its corresponding DEA result to identify subtype-specific functional pathways[38]. To this end, a geneset background was pre-prepared which includes all gene sets derived from GO biological processes (`c5.bp.v7.1.symbols.gmt`) from The Molecular Signatures Database (MSigDB, https://www.gsea-msigdb.org/gsea/msigdb/index.jsp). You can download other interested background for your own study.

```
# MUST locate ABSOLUTE path of msigdb file
MSIGDB.FILE <-
```

```
  system.file("extdata", "c5.bp.v7.1.symbols.xls", package = "MOVICS", mustWork = TRUE)
```

Likewise, these identified specific pathways should pass the significance threshold (*e.g.*, nominal $P$ value < 0.05 and adjusted $P$ value < 0.25) and must not overlap with any pathways identified for other subtypes. After having the subtype-specific pathways, genes that are inside the pathways are retrieved to calculate a single sample enrichment score by using GSVA R package[39-42]. Subsequently, subtype-specific enrichment score will be represented by the mean or median value within the subtype, and will be further visualized by diagonal heatmap.

```
# run GSEA to identify up-regulated GO pathways using results from edgeR
gsea.up <-
  runGSEA(moic.res     = cmoic.brca,
          dea.method   = "edger", # name of DEA method
          prefix       = "TCGA-BRCA", # MUST be the same of argument in runDEA()
          dat.path     = getwd(), # path of DEA files
          res.path     = getwd(), # path to save GSEA files
          msigdb.path  = MSIGDB.FILE, # MUST be the ABSOLUTE path of msigdb file
          norm.expr    = fpkm, # use normalized expression to calculate enrichment score
          dirct        = "up", # direction of dysregulation in pathway
          p.cutoff     = 0.05, # p cutoff to identify significant pathways
          p.adj.cutoff = 0.25, # padj cutoff to identify significant pathways
          gsva.method  = "gsva", # method to calculate single sample enrichment score
          norm.method  = "mean", # method to calculate subtype-specific enrichment score
          fig.name     = "UPREGULATED PATHWAY HEATMAP")
#> --all samples matched.
#> GSEA done...
#> --log2 transformation done for expression data.
#> Estimating GSVA scores for 50 gene sets.
#> Estimating ECDFs with Gaussian kernels
#>   |                                                                              |
```

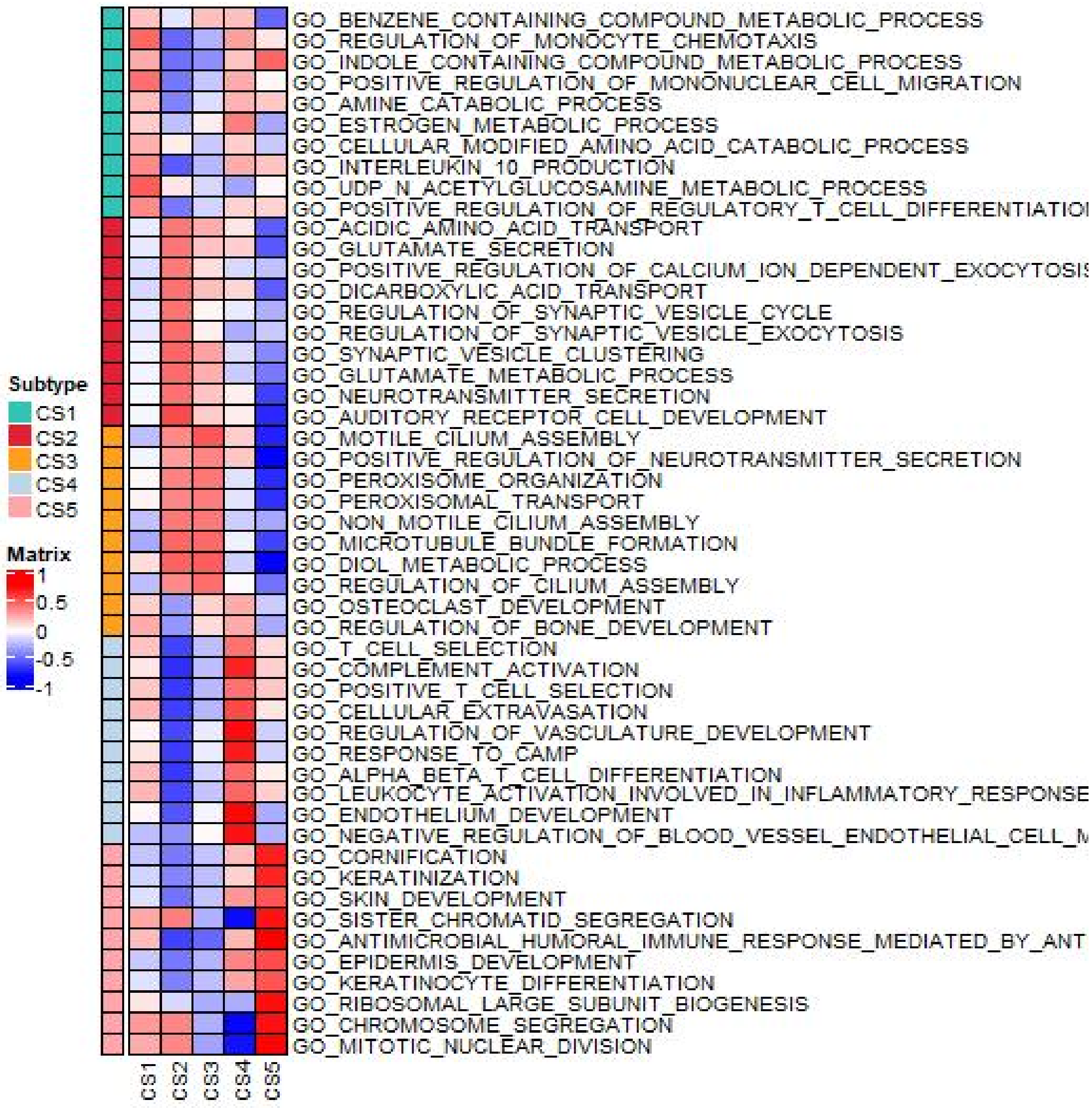

Figure 14: Heatmap of subtype-specific upregulated pathways using edgeR algorithm for 5 identified subtypes in TCGA-BRCA cohort.

Check some columns of GSEA results for the first subtype (CS1).

```
print(gsea.up$gsea.list$CS1[1:6,3:6])
```

Table 8: Demo of GSEA results for the first cancer subtype (CS1) of breast cancer in TCGA-BRCA cohort.

| | setSize | enrichmentScore | NES | pvalue |
|---|---|---|---|---|
| GO_EMBRYONIC_MORPHOGENESIS | 459 | -0.4531629 | -1.465422 | 0.001002 |
| GO_SYNAPTIC_SIGNALING | 441 | -0.5153428 | -1.663946 | 0.001003 |
| GO_REGULATION_OF_ION_TRANSPORT | 420 | -0.4868800 | -1.572493 | 0.001004 |
| GO_EPITHELIAL_CELL_DIFFERENTIATION | 470 | -0.4801548 | -1.552894 | 0.001005 |
| GO_AXON_DEVELOPMENT | 403 | -0.4788922 | -1.543713 | 0.001006 |
| GO_REGULATION_OF_TRANS_SYNAPTIC_SIGNALING | 293 | -0.5082236 | -1.626776 | 0.001006 |

Also check results of subtype-specific enrichment scores.

```
head(round(gsea.up$grouped.es,3))
```

Then try results derived from DESeq2 (or limma if you want) to identify subtype-specific downregulated pathways.

Table 9: Demo of subtype-specific enrichment scores among 5 identified subtypes of breast cancer in TCGA-BRCA cohort.

| | CS1 | CS2 | CS3 | CS4 | CS5 |
|---|---|---|---|---|---|
| GO_BENZENE_CONTAINING_COMPOUND_METABOLIC_PROCESS | 0.148 | -0.118 | 0.156 | 0.160 | -0.503 |
| GO_REGULATION_OF_MONOCYTE_CHEMOTAXIS | 0.425 | -0.500 | -0.274 | 0.256 | 0.040 |
| GO_INDOLE_CONTAINING_COMPOUND_METABOLIC_PROCESS | 0.219 | -0.472 | -0.382 | 0.142 | 0.430 |
| GO_POSITIVE_REGULATION_OF_MONONUCLEAR_CELL_MIGRATION | 0.401 | -0.453 | -0.212 | 0.220 | -0.012 |
| GO_AMINE_CATABOLIC_PROCESS | 0.172 | -0.419 | -0.140 | 0.196 | 0.132 |
| GO_ESTROGEN_METABOLIC_PROCESS | 0.119 | -0.231 | 0.013 | 0.356 | -0.305 |

```r
# run GSEA to identify down-regulated GO pathways using results from DESeq2
gsea.dn <- runGSEA(moic.res     = cmoic.brca,
                   dea.method   = "deseq2",
                   prefix       = "TCGA-BRCA",
                   msigdb.path  = MSIGDB.FILE,
                   norm.expr    = fpkm,
                   dirct        = "down",
                   p.cutoff     = 0.05,
                   p.adj.cutoff = 0.25,
                   gsva.method  = "ssgsea", # switch to ssgsea
                   norm.method  = "median", # switch to median
                   fig.name     = "DOWNREGULATED PATHWAY HEATMAP")
#> --all samples matched.
#> GSEA done...
#> --log2 transformation done for expression data.
#> Estimating ssGSEA scores for 50 gene sets.
#>   |                                                                               |
```

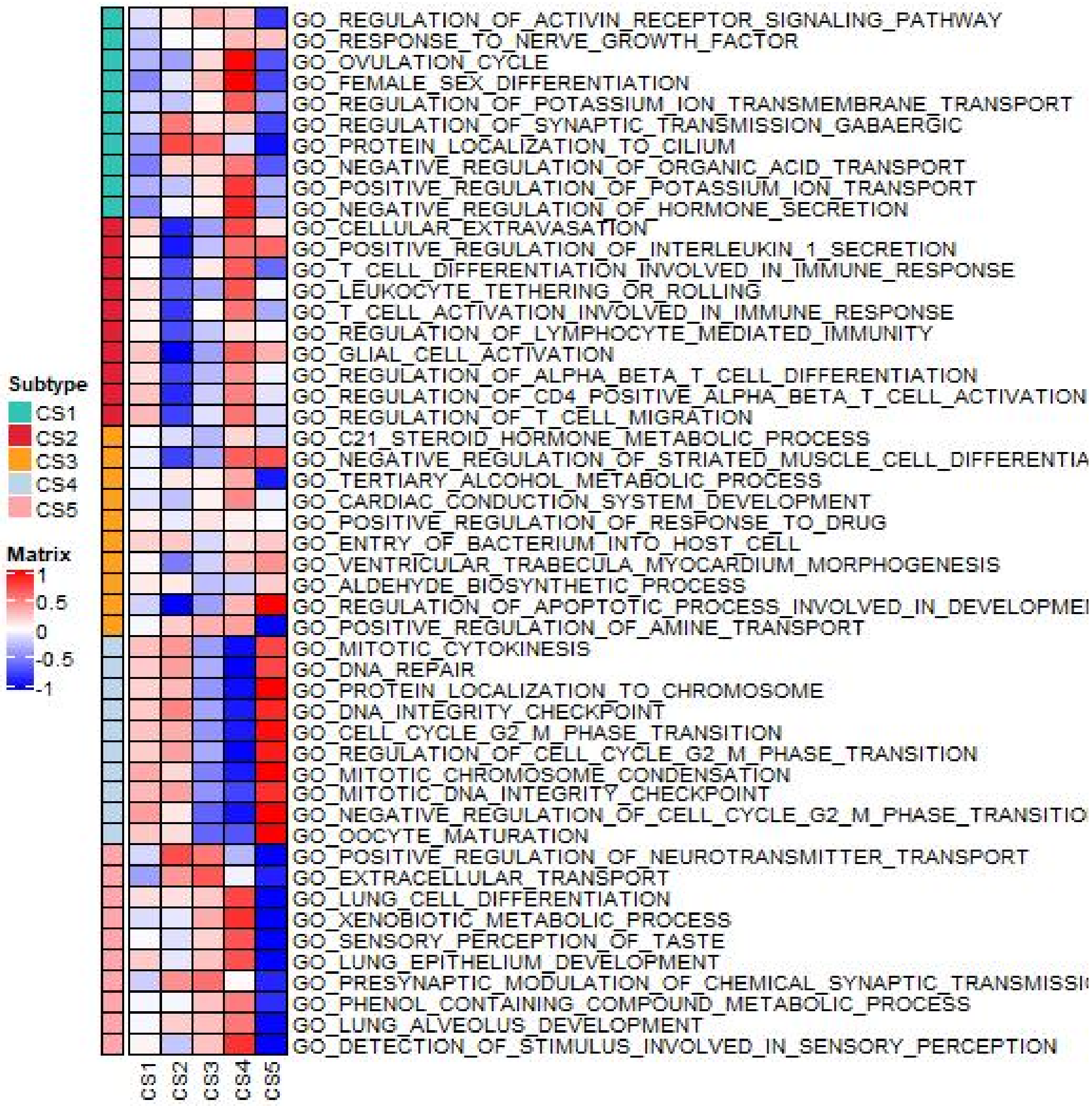

Figure 15: Heatmap of subtype-specific downregulated pathways using limma algorithm for 5 identified subtypes in TCGA-BRCA cohort.

**4.2.3.4 run nearest template prediction in external cohort** In this part, our core purpose is to predict the possible subtypes of each sample in the external dataset of Yau cohort. MOVICS harnesses nearest template prediction (NTP) which can be flexibly applied to cross-platform, cross-species, and multiclass predictions without any optimization of analysis parameters[43,44]. Only one thing have to do is generating a template file, which has been fortunately prepared already.

```
# run NTP in Yau cohort by using up-regulated biomarkers
brca.pred <-
  runNTP(expr      = brca.yau$mRNA.expr,
         templates = marker.up$templates, # template has been prepared in runMarker()
         scale     = TRUE, # scale input data
         center    = TRUE, # center input data
         doPlot    = TRUE, # to generate heatmap
         fig.name  = "NTP HEATMAP FOR YAU")
#> --original template has 500 biomarkers and 277 are matched in external expression profiling.
#> cosine correlation distance
#> 682 samples; 5 classes; 41-66 features/class
```

```
#> serial processing; 1000 permutation(s)...
#> predicted samples/class (FDR<0.05)
```

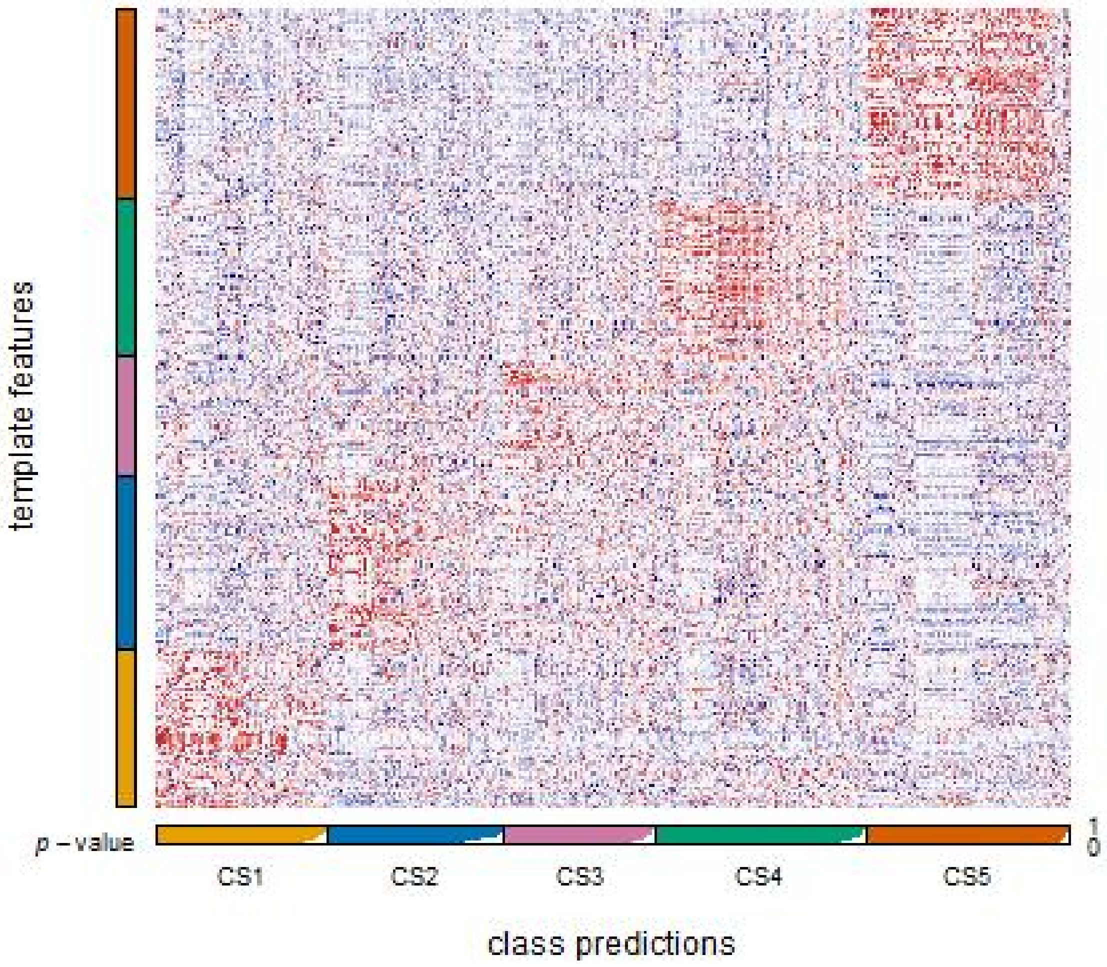

Figure 16: Heatmap of NTP in Yau cohort using subtype-specific upregulated biomarkers identified from TCGA-BRCA cohort

```
#>
#>  CS1  CS2  CS3  CS4  CS5 <NA>
#>  101   93   80  119  139  150
```

```
head(brca.pred$ntp.res)
```

Table 10: Demo of predicted subtypes in Yau cohort by NTP using subtype-specific upregulated biomarkers identified from TCGA-BRCA cohort.

| | prediction | d.CS1 | d.CS2 | d.CS3 | d.CS4 | d.CS5 | p.value | FDR |
|---|---|---|---|---|---|---|---|---|
| 107 | CS2 | 0.7245 | 0.5272 | 0.7355 | 0.7481 | 0.7512 | 0.001 | 0.0019 |
| 109 | CS1 | 0.5546 | 0.7607 | 0.7217 | 0.7329 | 0.7288 | 0.001 | 0.0019 |
| 11 | CS1 | 0.6590 | 0.7131 | 0.7557 | 0.7564 | 0.7716 | 0.002 | 0.0034 |
| 110 | CS2 | 0.7497 | 0.5327 | 0.7766 | 0.7669 | 0.7569 | 0.001 | 0.0019 |
| 111 | CS1 | 0.6852 | 0.7181 | 0.7167 | 0.7913 | 0.7909 | 0.013 | 0.0186 |
| 113 | CS4 | 0.6814 | 0.7505 | 0.7429 | 0.6440 | 0.7306 | 0.007 | 0.0107 |

Above an object `brca.pred` is prepared which is similar in structure of object returned by `getMOIC()` but only stores `clust.res` which can be passed to functions within **COMP Module** if additional data is available. For example, we can compare the survival outcome of the predicted 5 cancer subtypes in Yau cohort.

```
# compare survival outcome in Yau cohort
surv.yau <-
  compSurv(moic.res         = brca.pred,
           surv.info        = brca.yau$clin.info,
           convt.time       = "m", # switch to year
           surv.median.line = "hv", # switch to both
           fig.name         = "KAPLAN-MEIER CURVE OF NTP FOR YAU")
#> --a total of 682 samples are identified.
#> --cut survival curve up to 10 years.
```

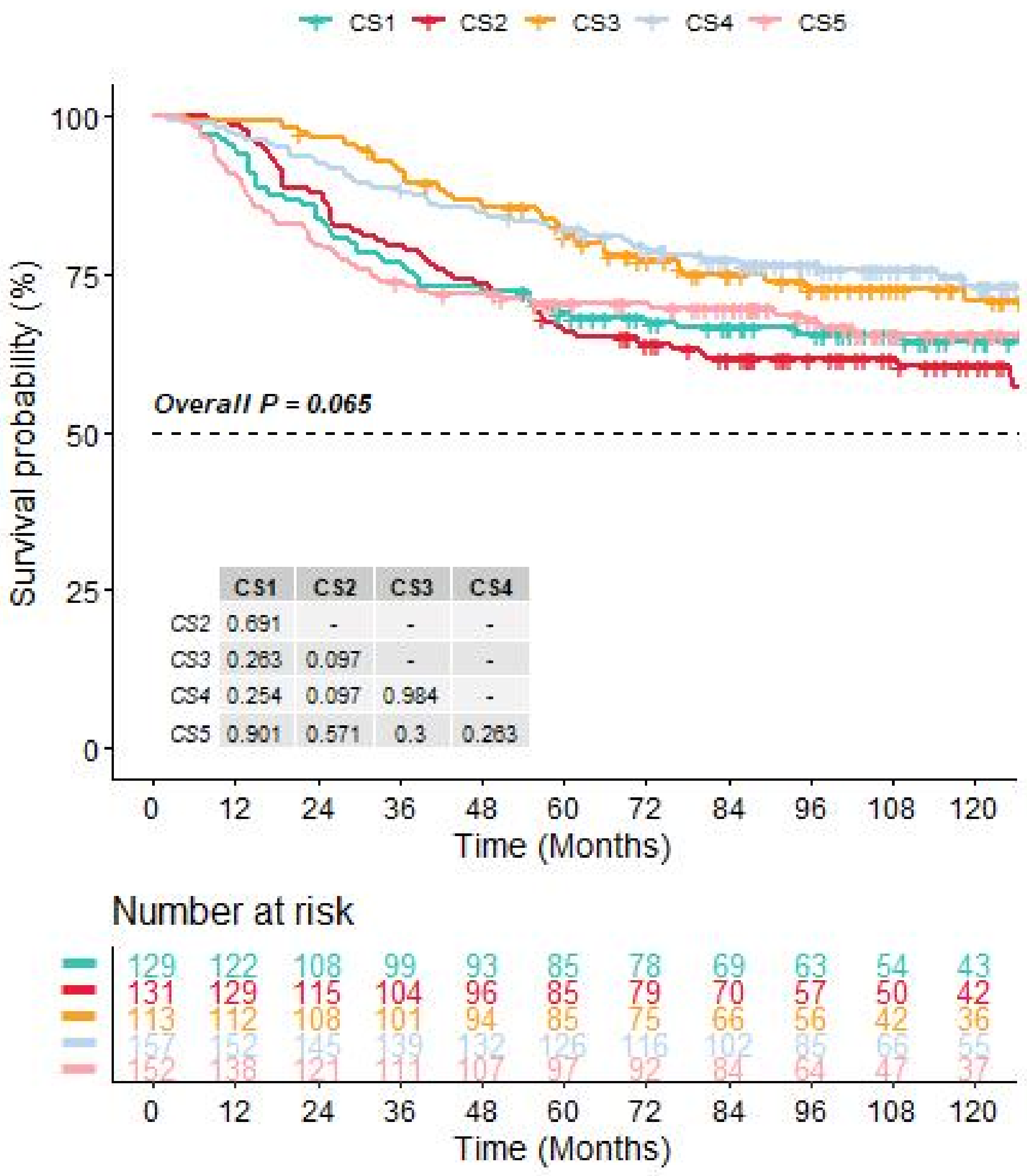

Figure 17: Kaplan-Meier survival curve of predicted 5 subtypes of breast cancer in Yau cohort.

```r
print(surv.yau)
#> $fitd
#> Call:
#> survdiff(formula = Surv(futime, fustat) ~ Subtype, data = mosurv.res,
#>     na.action = na.exclude)
#>
#>                N Observed Expected (O-E)^2/E (O-E)^2/V
#> Subtype=CS1 129       47     41.8     0.644     0.791
#> Subtype=CS2 131       55     42.8     3.445     4.259
#> Subtype=CS3 113       32     40.5     1.794     2.187
#> Subtype=CS4 157       44     56.3     2.676     3.570
#> Subtype=CS5 152       50     46.5     0.257     0.324
#>
#>  Chisq= 8.8  on 4 degrees of freedom, p= 0.07
#>
#> $fit
#> Call: survfit(formula = Surv(futime, fustat) ~ Subtype, data = mosurv.res,
#>     na.action = na.exclude, error = "greenwood", type = "kaplan-meier",
#>     conf.type = "plain")
#>
#>       n events median 0.95LCL 0.95UCL
#> CS1 129     47     NA      NA      NA
#> CS2 131     55    205     125      NA
#> CS3 113     32    236      NA      NA
#> CS4 157     44    222     191      NA
#> CS5 152     50     NA      NA      NA
#>
#> $overall.p
#> [1] 0.06513969
#>
#> $pairwise.p
#>
#>  Pairwise comparisons using Log-Rank test
#>
#> data:  mosurv.res and Subtype
#>
#>     CS1   CS2   CS3   CS4
#> CS2 0.691 -     -     -
#> CS3 0.263 0.097 -     -
#> CS4 0.254 0.097 0.984 -
#> CS5 0.901 0.571 0.300 0.263
#>
#> P value adjustment method: BH
```

We can further check the agreement between the predicted subtype and PAM50 classification.

```r
# compare agreement in Yau cohort
agree.yau <-
  compAgree(moic.res  = brca.pred,
            subt2comp = brca.yau$clin.info[, "PAM50", drop = FALSE],
            doPlot    = TRUE,
            fig.name  = "YAU PREDICTEDMOIC WITH PAM50")
#> --all samples matched.
```

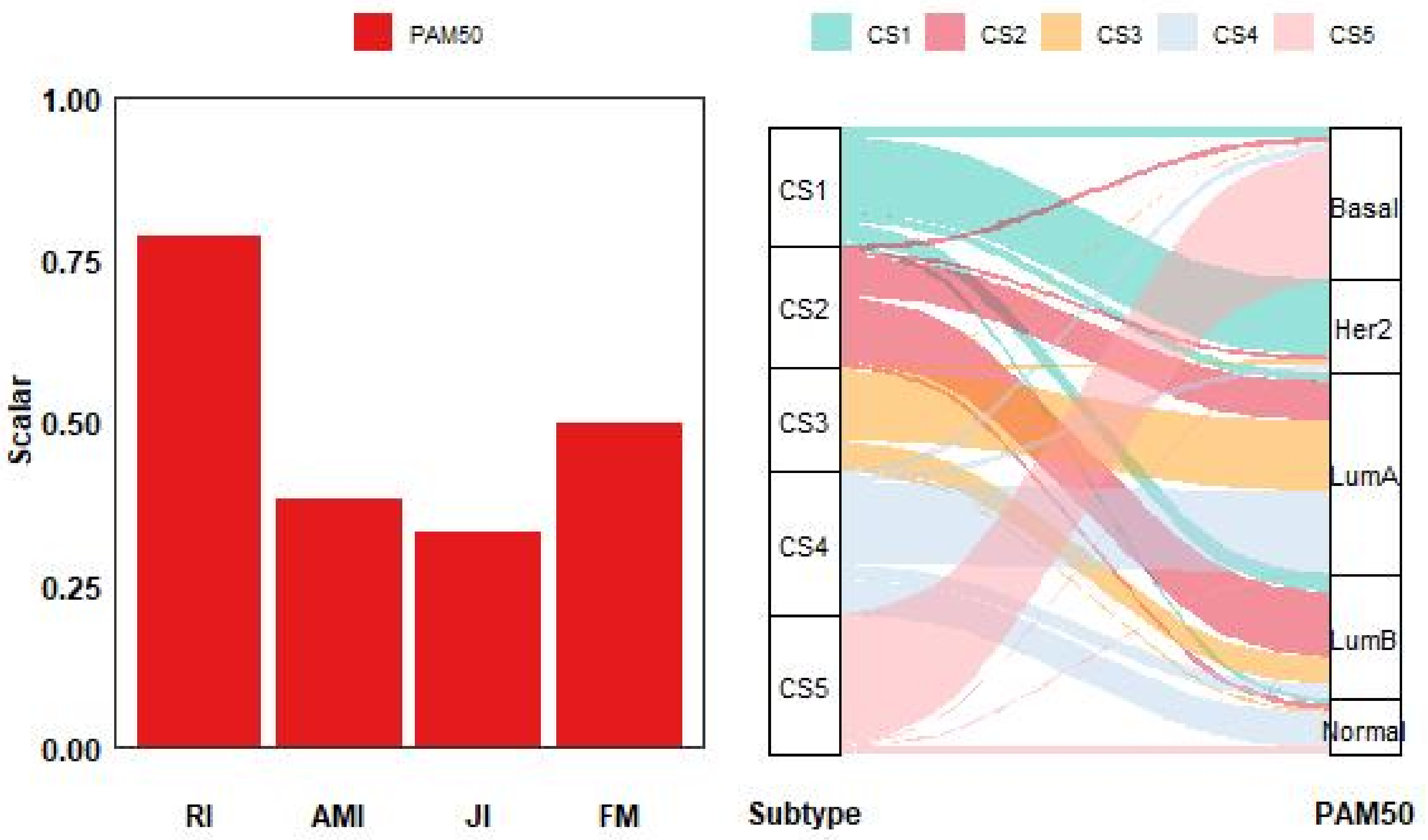

Figure 18: Agreement of predicted 5 subtypes of breast cancer with PAM50 classification in Yau cohort.

```
print(agree.yau)
```

Table 11: Agreement of 5 predicted subtypes of breast cancer with PAM50 classification in Yau cohort.

| current.subtype | other.subtype | RI | AMI | JI | FM |
|---|---|---|---|---|---|
| Subtype | PAM50 | 0.7828835 | 0.3826727 | 0.3313396 | 0.4988811 |

It is clearly that the predicted subtypes of breast cancer in Yau cohort can distinguish prognosis well and also show similar pattern of agreement with PAM50 classification as compared to TCGA-BRCA to some extent.

#### 5 Little Trick

Indeed, the `moic.res` parameter is necessary for almost all the functions, especially for downstream analyses. But in fact, you can fool downstream modules with little tricks. The core information stored in `moic.res` is a data.frame named `clust.res` with a `samID` column (character) for samples name (should be exactly the same with row names) and a `clust` column (integer) for subtype indicator. Another information required for some functions in downstream analyses is `mo.method`, a string value that is usually considered as prefix for output files. For example, if I want to check the prognostic value of PAM50 classification, the only thing is to prepare is a pseudo `moic.res` object with a customized `mo.method` just like below:

```
# a list of original clinical information as `clust.res` and `mo.method`
pseudo.moic.res                 <- list("clust.res" = surv.info,
                                        "mo.method" = "PAM50")

# make pseudo samID
```

```r
pseudo.moic.res$clust.res$samID <- rownames(pseudo.moic.res$clust.res)

# make pseudo clust using a mapping relationship
pseudo.moic.res$clust.res$clust <- sapply(pseudo.moic.res$clust.res$PAM50,
                                          switch,
                                          "Basal"   = 1, # relabel Basal as 1
                                          "Her2"    = 2, # relabel Her2 as 2
                                          "LumA"    = 3, # relabel LumA as 3
                                          "LumB"    = 4, # relabel LumnB as 4
                                          "Normal"  = 5) # relabel Normal as 5
```

Check how the pseudo `moic.res` object looks like.

```r
head(pseudo.moic.res$clust.res)
#>                 fustat futime PAM50 pstage age        samID clust
#> BRCA-A03L-01A      0   2442  LumA     T3  34 BRCA-A03L-01A     3
#> BRCA-A04R-01A      0   2499  LumB     T1  36 BRCA-A04R-01A     4
#> BRCA-A075-01A      0    518  LumB     T2  42 BRCA-A075-01A     4
#> BRCA-A08O-01A      0    943  LumA     T2  45 BRCA-A08O-01A     3
#> BRCA-A0A6-01A      0    640  LumA     T2  64 BRCA-A0A6-01A     3
#> BRCA-A0AD-01A      0   1157  LumA     T1  83 BRCA-A0AD-01A     3
```

Just keep in mind how your interested subtypes are mapped and feel free to use such pseudo object to perform downstream analyses such as `compSurv()` as follows:

```r
# survival comparison
pam50.brca <-
  compSurv(moic.res         = pseudo.moic.res,
           surv.info        = surv.info,
           convt.time       = "y", # convert day unit to year
           surv.median.line = "h", # draw horizontal line at median survival
           fig.name         = "KAPLAN-MEIER CURVE OF PAM50 BY PSEUDO")
#> --a total of 643 samples are identified.
#> --removed missing values.
#> --leaving 642 observations.
#> --cut survival curve up to 10 years.
```

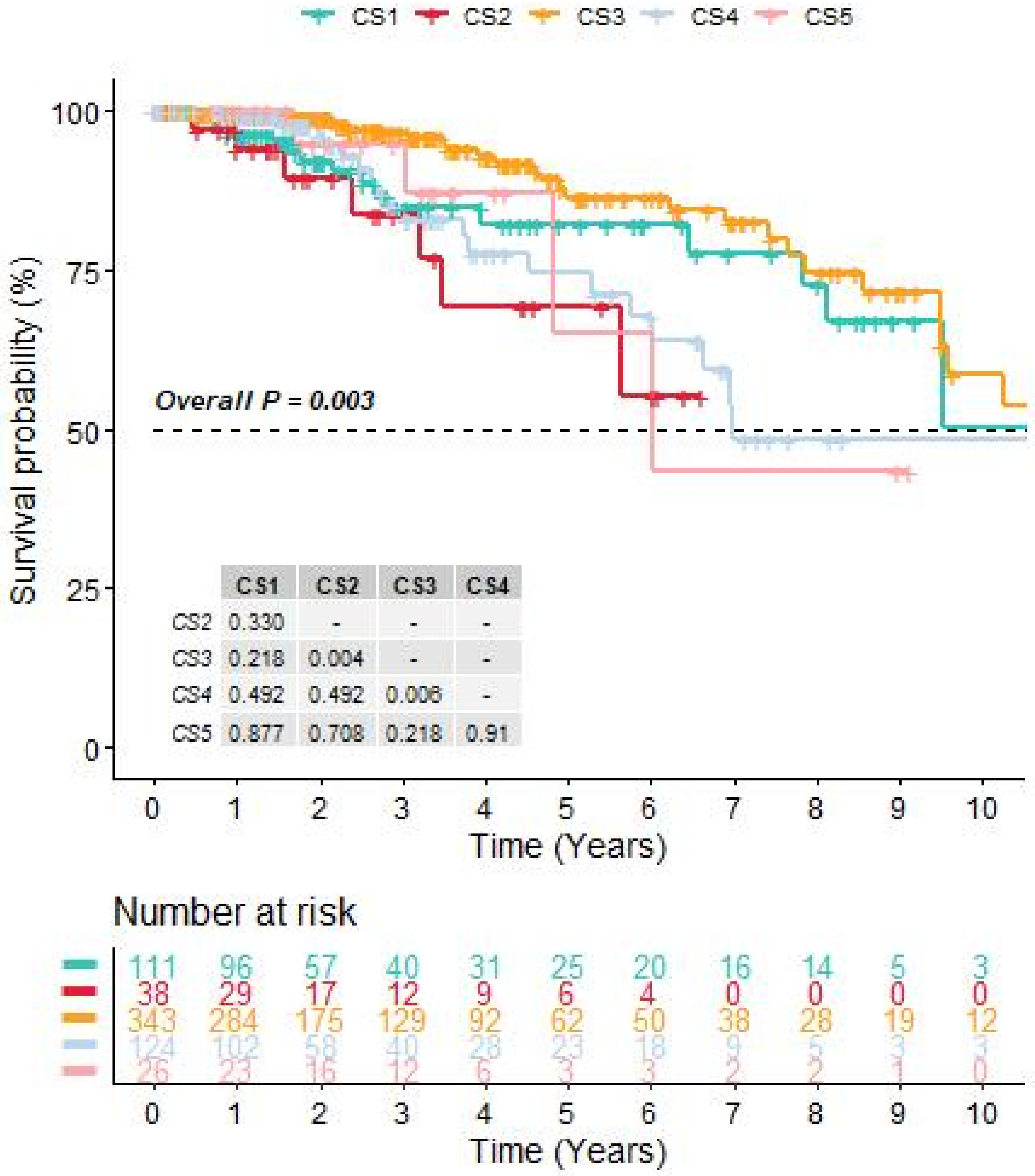

Figure 19: Kaplan-Meier survival curve of PAM50 subtypes of breast cancer with pseudo input in TCGA-BRCA cohort.

#### 6 Summary

In summary, MOVICS provides a suite for multi-omics integration and visualization in cancer subtyping.
